# Genetic and functional dissection of the glutamate-proline pathway reveals a shortcut for glutamate catabolism in *Leishmania*

**DOI:** 10.1101/2024.02.06.579205

**Authors:** Gustavo Daniel Campagnaro, Angela Kaysel Cruz

## Abstract

Trypanosomatids are early-divergent eukaryotes that have adapted to parasitism. During their life cycles, these parasites switch between a mammalian and an invertebrate host, and the ability to adapt their metabolism to different nutritional sources is detrimental for their success. In the invertebrate host, these protists have access to high amounts of amino acids and efficiently utilise it for energy production. Proline is a particularly efficient energy source for trypanosomes. Glutamate is also efficiently used by *Trypanosoma cruzi*, but it needs to be converted into proline prior to its catabolism. By employing a series of genetic modifications and functional analysis, we show here that *Leishmania* parasites, the causative agents of leishmaniases, can utilise proline, glutamate and glutamine as energy sources, and although these parasites possess all the genes necessary for the biosynthesis of proline from glutamate, this pathway has, at best, limited function, with at least one of its components (pyrroline-5-carboxylate reductase) assuming divergent functions in different life cycle stages of the parasite. In fact, we show that the catabolism of glutamate is independent of proline biosynthesis and the former is most likely directly imported into the mitochondrion and catabolised to recover the cellular redox metabolism and increase mitochondrial membrane potential. Moreover, our data suggest a relevant role for glutamate dehydrogenase in nutritional stress response in *Leishmania*. These findings highlight relevant differences in amino acid metabolism between *Trypanosoma* and *Leishmania* and suggest a diversification in amino acid metabolic pathways within Trypanosomatidae.

## 1. Introduction

Glutamate (Glu) is a non-essential amino acid that can be considered a metabolic hub, as it can be generated from and converted to a wide variety of metabolites. Remarkably, Glu is the most abundant intracellular amino acid and constitutes an important Nitrogen donor for the biosynthesis of other biomolecules, particularly of other amino acids: most of the carbons in arginine (Arg) and all the carbons present in proline (Pro) are derived from Glu. Glu can be obtained from the extracellular environment via active transport, as well as by its biosynthesis from a wide diversity of metabolic pathways (1). Regarding the connection between amino acid metabolic pathways, Glu can be synthesised from alanine, arginine, aspartate, glutamine (Gln), histidine, and proline via multi-enzymatic steps (1).

Besides its primary role as a constituent of proteins, Glu constitutes a valuable energy source as it can be catabolized into α-ketoglutarate, a tricarboxylic acid (TCA) cycle intermediate, via three different pathways: (1) deamination by glutamate dehydrogenase (GluDH), which is the main route for Glu catabolism and generates α-ketoglutarate and ammonia, (2) transference of the amino group from glutamate to pyruvate by either alanine aminotransferase (ALAT) or tyrosine aminotransferase (TAT), with the production of alanine and α-ketoglutarate, and (3) the conjugation of glutamate with oxaloacetate by aspartate aminotransferase (ASAT) to generate α-ketoglutarate and aspartate (2–4).

Importantly, in most organisms, GluDH is a mitochondrial enzyme, and Glu can reach the mitochondrial matrix via (1) active transport from the cytosol (5, 6) or, most commonly, (2) as part of the redox shuttle that occurs in the Glu-Pro pathway (7, 8). In most organisms, in the Glu-Pro pathway, Glu is phosphorylated into γ-glutamyl phosphate by γ-glutamyl kinase (GK) at the expense of one ATP, then γ-glutamyl phosphate is reduced to glutamate-γ-semialdehyde (GSA) by γ-glutamyl phosphate reductase (GPR) with the oxidation of one NADPH. In an aqueous environment, GSA is spontaneously converted into pyrroline-5-carboxylate (P5C), which is then further reduced into Pro by P5C reductase (P5CR) with the expense of a NAD(P)H. Pro is then transported into the mitochondria and catabolised into P5C by Pro dehydrogenase (ProDH) with the reduction of FAD^-^ to FADH_2_. Next, P5C is oxidized back to Glu by P5CDH with the reduction of NAD^+^ to NADH (7–9). The FADH_2_ and NADH generated in this pathway feed the respiratory chain with electrons and propel energy production. Importantly, a shorter P5C-Pro cycle can be established if P5C accumulates at high levels within the mitochondria (10). Furthermore, the physical separation of the Glu-Pro pathway steps between cytosol and mitochondria is important for balancing cellular redox state and the ratios of NAD(P)^+^ and NAD(P)H_2_, implying a connection with other metabolic pathways, such as glycolysis and the pentose phosphate pathway (7, 11).

Pro constitutes a major energy source utilised by trypanosomatids a group of early-divergent eukaryotic parasitic protozoa (12), within the insect vector and is the preferred energy source in the absence of glucose *in vitro* (13–15). *Trypanosoma brucei*, the causative agent of sleeping sickness, is auxotrophic for Pro, which is the only amino acid able to sustain the growth in the absence of glucose (14, 15). *Trypanosoma cruzi*, the causative agent of Chagas’ disease, in addition to Pro uptake from the host, displays a functional Glu-Pro pathway and, in fact, it is currently accepted that the use of Glu as energy source in *T. cruzi* is highly dependent on the biosynthesis of Pro (4, 10, 16). *Leishmania* spp., which causes leishmaniases, can take Pro up from the host and also possess all genes necessary for a functional Glu-Pro pathway, although very little is known about Pro biosynthesis in this genus (4, 17, 18).

It has been shown that Pro, as well as Gln, cysteine, and tyrosine, but not Glu, are necessary for continuous *in vitro* growth of *Leishmania braziliensis* and *Leishmania donovani* (19). Glu, Gln and Pro are, however, non-essential for *Leishmania mexicana* growth (20), which would imply the existence of a functional (and efficient) Glu-Pro pathway in this species that even allows the secretion of Pro into the culture medium (21). Metabolomic data from *L. mexicana* promastigotes, however, show that the taken up Glu is mostly directed to fuelling the TCA cycle and only a minute proportion is used for Pro biosynthesis (22–24), which could suggest an allosteric regulation of this pathway when Pro can be scavenged from the environment in sufficient amounts (24, 25), but also reveal an important role for Glu in *Leishmania* energy metabolism. In amastigote forms, central Carbon metabolism of *Leishmania* is rewired and glucose and fatty acids become the preferential energy sources (23, 26, 27), although glucose-deprived amastigotes can also utilise amino acids for energy production (22, 27).

To investigate the use of Glu as an energy source and the functionality of the Glu-Pro pathway in *Leishmania*, we employed CRISPR/Cas9 to knockout members of the Glu-Pro pathway in *Leishmania braziliensis* and investigated the ability of the mutant cell lines to recover from nutritional stress utilizing glucose or amino acids as single Carbon sources, which revealed that although glucose and Pro are preferred, Glu is also an efficient energy sources. Importantly, cells knocked out for ProDH became unable to utilize Pro, but could recover their cellular metabolism when provided with Glu. The knockout of GluDH1 completely abolished the functionality of the Glu-Pro pathway in restoring energy metabolism. Moreover, we show that when the catabolism of Pro is blocked, Glu itself is used to fuel mitochondrial metabolism, and increase mitochondrial membrane potential (ΔΨ_m_). Together, our results suggest the canonical Glu-Pro pathway may have a very limited functionality in *L. braziliensis* and, different from trypanosomes, that Glu can be directly utilised as an energy source.

## 2. Materials and Methods

### 2.1. Parasite culture and genetic manipulation

*Leishmania amazonensis* (MHOM/BR/73/M2269), *Leishmania major* (MRHO/SU/59/P) and *Leishmania braziliensis* (MHOM/BR/75/M2903) wildtype strains were utilised. *L. braziliensis* expressing T7 RNAP, Cas9 and TdTomato, here referred to as pT007, were used for genetic manipulation using conditions previously established in our laboratory (28). All genetic modifications were verified by PCR using primers listed in Supplementary Table 1. Characteristic PCR results for each cell line generated are shown in Supplementary Figure 1.

All parasite cell lines were maintained as promastigotes at 26°C in M199 culture medium (Sigma-Aldrich M0393) supplemented with 10% fetal bovine serum (FBS), 100 U/ml penicillin, 100 μg/ml streptomycin, 2 mM L-glutamine, 40 mM Hepes, 0.1 mM adenine (in 50 mM Hepes), 5mg/mL hemin (in 50% triethanolamine), and 1 mg/ml 6-biotin. Media for maintenance of *L. braziliensis* was further supplemented with 1µM biopterin.

For differentiation into axenic amastigotes, metacyclic forms of *L. braziliensis* were enriched using Ficoll gradient from cultures in stationary phase. 2×10^6^ metacyclic forms were seeded into 10 mL of FBS and transferred to humid atmosphere at 33°C and 5% CO_2_. From the fourth day of differentiation, cells presented smaller size, rounded shape, and limited movement, and were considered amastigotes (28, 29).

### 2.2. Nutritional stress and recovery

Cells in logarithmic growth were harvested by centrifugation (10 min at 1,300×g), washed twice in PBS and subjected to nutritional stress by incubation in PBS for established periods: two hours for *L. amazonensis* and *L. major*, and four hours for *L. braziliensis*, unless stated otherwise. As control, non-stressed samples that were always incubated in M199 were run alongside stressed samples.

After, cells were transferred to M199, PBS, or PBS containing 3 mM of Pro, Glu, Gln or Arg, and allowed to recover for four hours. In order to avoid disproportionate growth in some samples, and consequent skewing of results, M199 without FBS was used in these assays. Cells kept in M199 without FBS (ie, not starved in PBS) for the course of the assay were considered as a non-stressed control.

Next, 0.4mg of MTT (M5655; Sigma-Aldrich) was added to each well of a 96-well plate, and incubated for 24h at 26°C. The reduced MTT crystals were collected by centrifugation at 3000×g for 10 min, and the supernatant was discarded by inversion. MTT crystals were solubilised by addition of 100 µL of DMSO to each well, and plates were read at 570 nm in SpectraMax i3 (Molecular Devices).

### 2.3. Amino acid dosage

The measurement of intracellular proline was carried out as previously described (10, 30) using 3×10^7^ cells per sample. Briefly, *L. braziliensis* wildtype cells were collected by centrifugation (1,300 × *g* for 5 min), washed twice with PBS and transferred to FBS-free M199 (non-stressed) or to PBS (nutritional stress) for 2h. After that, cells were pelleted, washed once with PBS and resuspended in appropriate buffer as reported by Marchese et al. (2020), and let to recover for two hours. At the time of sample collection, the parasites were pelleted by centrifugation at 4°C, washed twice with cold PBS, resuspended in lysis buffer (100 mM Tris-HCl pH 8,1; 250 mM sorbitol, 1 mM EDTA, 1% Triton X-100 (v/v), 1 mM phenylmethylsulfonyl fluoride (PMSF), and 2× cOmplete proteinase inhibitor cocktail (Roche)), and submitted to two cycles of snap freezing in liquid N_2_ and thawing. Crude extracts were clarified by centrifugation (15,000 × *g* for 15 min at 4°C). 100 µL of supernatant was removed and mixed with equal volume of 20% (w/v) trichloroacetic acid for 1h in ice for deproteinization. Proteins were separated by centrifugation (20,000 × *g* for 30 min at 4 °C), and 200 µL of the resultant supernatant was used for Pro quantification following a ninhydrin assay (31).

Two hundred µL of sample extract were mixed with 200 μl of glacial acetic acid and 200 μL of fresh ninhydrin solution (250 mg of ninhydrin diluted in a mixture of 6 mL of glacial acetic acid and 4 mL of 6M H_3_PO_4_) in a cryovial and boiled for 1h. Samples were transferred to ice and 400 μL of toluene were added to each sample. Samples were mixed by vortexing for 20 seconds and transferred back to ice for 10 min. 100 μL of the red-coloured organic phase was collected, transferred to a 96-well polypropylene plate and read at 520 nm in SpectraMax i3 (Molecular Devices).

The amount of intracellular Glu was measured in 1×10^6^ cells using the kit Glutamate-Glo (J7021; Promega) according to the manufacturer’s instructions. For that, pT007 and *Lbr*GluDH1-KO cells were collected by centrifugation (1,300 × *g* for 5 min), washed twice with PBS and transferred either to FBS-free M199 (non-stressed) or to PBS (nutritional stress) for one hour. After that, cells were pelleted, washed once with PBS and resuspended in appropriate buffer: cells from the non-stressed group were resuspended back into FBS-free M199 and the cells starved in PBS were resuspended in PBS only or PBS containing 3mM of Pro, Glu or Gln. Samples were further incubated for one hour and then collected by centrifugation at 4 °C, washed three times with cold PBS and lysed with 0.6N HCl for five min on ice, which was neutralised by addition of an equal volume of 1M Tris base. Samples were distributed in black-walled flat-bottom 96-well plates and luminescence was recorded in Synergy2 (BioTek).

### 2.4. RT-qPCR

In order to measure transcript abundance across the life cycle of *L. braziliensis*, total RNA was extracted from procyclic forms collected from logarithmic growing cultures, as well as from metacyclic-enriched fractions, and axenic amastigote forms of pT007 cells. All RT-qPCR reactions were performed in the platform abi 7500 (Thermo Fisher Scientific) using Mix PowerUp SYBR Green (Thermo Fisher Scientific) and primers listed in Supplementary Table 1.

### 2.5. Western blotting

Protein expression in different life cycle forms was measured by western blotting. Procyclic, metacyclic and amastigote forms of cells expressing myc-tagged GK, GPR and P5CR were collected by centrifugation, washed two to three times in PBS (1,300×g for 10 min) at 4°C, resuspended in extraction buffer (SDS 2%, 50 mM Tris-Cl pH 7.4, 1 mM PMSF, 2x cOmplete protein inhibitor cocktail (Roche)), and boiled for 10 min. Protein concentrations were measured on a Nanodrop One instrument (Thermo Scientific). 30 µg of each sample were mixed with 0.2 V of 6x sample buffer (350 mM Tris-Cl pH 6.8, 30% glycerol, 10% SDS, 0.12 mg/mL bromophenol blue, 6% 2-mercaptoethanol) and boiled for three more minutes prior application onto the gel.

After transference, nitrocellulose membranes were blocked for >1h with 5% skimmed milk at room temperature and incubated overnight with primary anti-myc antibody (CST 2276S) at 1:10000 in 5% milk at 4°C. After, membranes were washed and incubated with secondary anti-mouse peroxidase-conjugated antibodies (GE Healthcare NA931V) for 1h. The membranes were revealed using the ECL kit (GE Healthcare RPN2232), and images were captured on ImageQuant LAS 4000 (GE Healthcare) device. All band density quantifications were performed in non-saturated images using ImageJ.

### 2.6. Immunofluorescence

*Lbr*GluDH1-myc cells were collected in exponential growth phase, washed once with PBS, resuspended in fresh complete M199 and incubated with 100 nM of MitoTracker DeepRed (MTDR) (M22426; Thermo Scientific) for 30 min. After, cells were collected by centrifugation, washed thrice in PBS and fixed with 3% paraformaldehyde for 10 min under gentle agitation. Cells were then washed with PBS and the paraformaldehyde was neutralised by addition of a 0.1% glycine solution. All samples were kept at 4°C until processing.

Cells were placed in a poly-lysine-coated slide, permeabilised with 0.2% Tween20 in PBS for 10 min, and blocked with 0.2% Tween20, 1% BSA in PBS (blocking solution), followed by incubation with anti-myc primary antibody diluted at 1:2000 in blocking solution for two hours. A secondary anti-mouse conjugated to Alexa488 (Invitrogen A11001) diluted at 1:500 in blocking solution was added for 30 min. Finally, nuclei and kinetoplasts were stained using Hoechst (Invitrogen H3570) at 2 μg/mL in PBS for 15 min. Images were acquired with an Axio Observer combined with an LSM 780 confocal device (Carl Zeiss). Images were processed with Fiji ImageJ free software.

### 2.7. Immunoprecipitation

The immunoprecipitation followed a previously published protocol (28) with minor modifications. Briefly, 2×10^8^ procyclic on the second day of growth or fully differentiated axenic amastigote forms of the myc-*Lbr*P5CR line were collected by centrifugation (1,300 x *g* for 10 min) at 4°C, washed twice with cold PBS and resuspended in lysis buffer (10 mM Tris pH 7.5, 50 mM KCl, 2 mM MgCl_2_, PMSF 2 mM, 1 mM DTT, 1% Triton X-100, 10% glycerol, 2x cOmplete protease inhibitor (Roche)). Lysis was initiated by incubation at 4°C for 30 minutes under mild agitation. Then, the cells were subjected to mechanical disruption using serial passaging in 29G needles. Cell debris were removed by centrifugation (10,000 x *g* for 10 min at 4°C) and the supernatant was incubated with 30 μL of protein A/G-coated magnetic beads (Genscript L00277) for 1 hour at 4 °C to remove nonspecifically bound proteins. After, precleared supernantants were recovered and incubated with beads that had been coated with the anti-myc antibody (CST 2276S) for one hour at room temperature. After 4h of agitation at 4°C, beads were recovered by magnetism and washed twice with lysis buffer. Bound proteins were eluted by boiling the beads with 30 μL of sample buffer (117 mM Tris-Cl pH 6.8, 10% glycerol, 3.3% SDS, 0.04 mg/mL bromophenol blue, 2% 2-mercaptoethanol) at 6x for 10 min. A fraction of the eluted proteins was used to confirm the efficiency of the pulldown by Western blotting, and the remaining proteins were sent to the University of Laval for mass spectrometry (MS) identification.

MS data was analysed using Scaffold (version Scaffold_5.2.2, Proteome Software Inc., Portland/OR). Protein identification threshold was set at 99% and only peptides identified with ≥95% of confidence were considered in the analysis. All proteins identified in the control samples were excluded from the analysis. Subsequently, the proteins identified exclusively in procyclic or in amastigote forms were subjected to GO Term analysis for “Biological Process” and “Molecular Function” in TritrypDB (32).

### 2.8. Assessment of mitochondrial potential by MitoTracker

pT007 and *Lbr*ProDH-KO cells were collected by centrifugation (1,300 × *g* for 10 min), washed once in PBS and resuspended in FBS-free M199 (non-stressed) or in PBS and incubated at 26°C for two hours. After, cells were pelleted, washed once with PBS and resuspended as appropriate: non-stressed cells were resuspended back in FBS-free M199, while cells starved in PBS were resuspended in FBS-free M199, PBS or PBS containing 3mM of Pro or Glu and let to recover for two hours. 30 min prior to sample collection, 100 nM of freshly diluted MTDR was added to each sample. When necessary, 10µM of Carbonyl cyanide 4-(trifluoromethoxy)phenylhydrazone (FCCP) (SML2959; Sigma-Aldrich), an uncoupler of mitochondrial oxidative phosphorylation, was added 30 min prior to the addition of MTDR. Cells were protected from light after the addition of either MTDR or FCCP. Each group contained 5×10^6^ cells.

At the time of collection, samples were transferred to ice for two minutes, and collected by centrifugation (1,300 × *g* for 5 min) at 4°C, washed twice with ice-cold PBS, and fixed with 3% paraformaldehyde in PBS for 10 min under gentle agitation, followed by a final wash with PBS. Cells were then resuspended in 1 mL of PBS and stored at 4°C until analysis. Unstained controls (which did not receive MTDR) were used to set the baseline of fluorescence and “dead” controls were added to confirm the accumulation of MTDR in actively respirating mitochondria (33, 34): for this, cells were fixed with 3% paraformaldehyde as described above, resuspended in FBS-free M199, and followed the same protocol as the “living” groups. All samples were filtered through 35 mm nylon mesh membranes and the quantification of MTDR was performed by flow-cytometry in a BD FACSCanto II or in a BD Accuri C6 Plus with similar outcomes. 30,000 events were collected per sample. The data was analysed and plotted using FlowJo v10 (https://www.flowjo.com). All data are plotted as Mean Fluorescence Intensity (MFI).

### 2.9. Statistical analysis

All data analyses were performed in GraphPad Prism 10.1.1. Analyses that compared several different conditions between two or more cell lines or treatment groups were performed using two-way ANOVA followed by Tukey’s post-test. Except where stated otherwise, comparisons made between more than two conditions for a single cell line or treatment group were analysed by one-way ANOVA followed by Tukey’s post-test. When only two conditions for the same cell line or two cell lines subjected to same condition were compared, unpaired two-tailed Student’s t-test was used.

### 2.10. Data Availability statement

The authors declare that all data supporting the findings of this study are presented within the paper and its supplementary files. All raw data are available from the corresponding authors upon request.

## Results

### *Leishmania* species efficiently utilise amino acids as energy sources and restore their redox potential after nutritional stress

Previous studies in *L. mexicana* have shown that amino acids constitute important energy sources for *Leishmania* promastigotes(22–24), and that different species consume different amounts of each amino acid from the culture medium (21). To test whether different species could use different amino acids as preferred energy sources, we subjected *L. amazonensis*, *L. major* and *L. braziliensis* promastigotes to 2 (for *L. amazonensis* and *L. major*) or 4 hours (for *L. braziliensis*) of nutritional stress in PBS and then recovery in PBS containing a single Carbon source for further 4h. The recovery of mitochondrial redox potential was assessed by incubation with MTT for 24h.

All species efficiently utilised glucose, Pro, Glu and Gln to restore their mitochondrial metabolism, with glucose and Pro being the preferred energy sources (Figure 1). Importantly, *L. braziliensis* utilised amino acids in a more efficient fashion than *L. amazonensis* and *L. major* and converted MTT at higher rates. Arginine, differently, could not be efficiently used to recover cell energy metabolism, likely because no known connection between arginine metabolism and energy metabolic pathways exists in trypanosomatids. In these organisms, Arginine is rather directed to polyamine biosynthesis (4, 35). Of note, the efficient recovery in cell energy metabolism was mirrored by a visual recovery in cell motility under the microscope.

**Figure 1.**
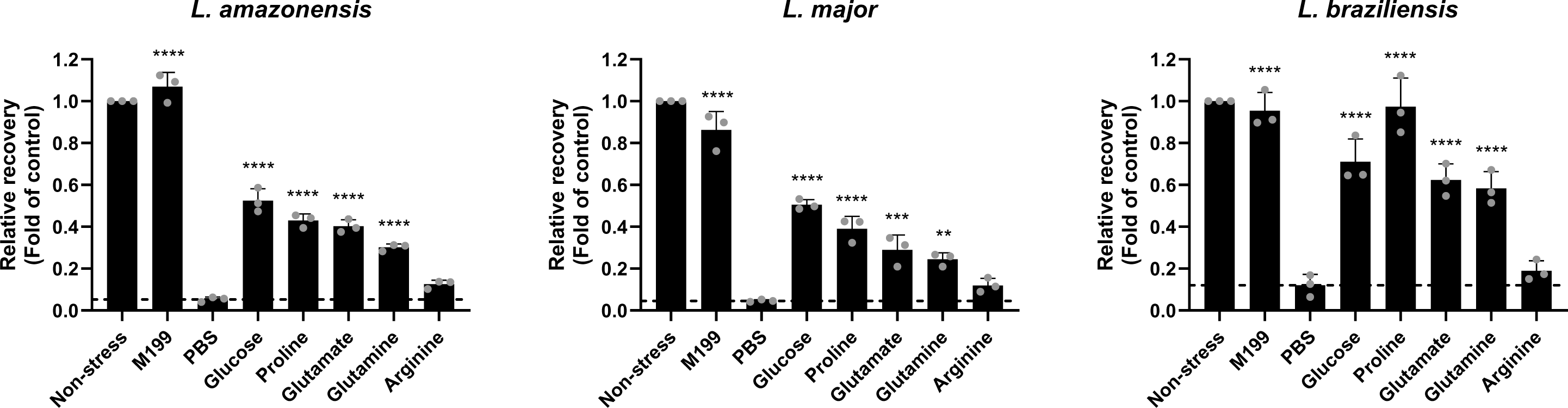
*Leishmania* species effectively utilise amino acids as energy sources. The ability of *L. amazonensis*, *L. major* and *L. braziliensis* wildtype cells subjected to stress in PBS and let to recover in the presence of a single nutritional source to recover their cellular metabolism and convert MTT was measured. Cells kept in M199 without serum were utilised as non-stressed controls. Statistical analysis was performed using One-way ANOVA followed by Dunnett’s post-test by comparing samples recovered in nutritious buffers to samples kept in PBS. **P<0.01; ***P<0.001; ****P<0.0001.

### *L. braziliensis* expresses enzymes involved in Pro biosynthesis *de novo* at different levels and these are not essential for the catabolism of amino acids in the recovery from nutritional stress

In trypanosomatids, the use of Glu and Gln as energy sources is believed to occur by the entry of Glu in the Glu-Pro pathway (4, 10, 16), which implicates in the *de novo* synthesis of Pro by the sequential activity of GK, GPR and P5CR before it is catabolysed to generate energy. Noteworthy, in more complex eukaryotes, the reactions catalysed by GK and GPR are coupled together by the action of a bifunctional enzyme, P5C synthetase. This also occurs in *T. cruzi*. Genes encoding for both GK and GPR were identified in *L. braziliensis* (LBRM2903_260032600 and LBRM2903_320042400, respectively), and the amino acid sequences of these proteins are closely related to the *Saccharomyces cerevisiae* GK and GPR (Figure 2A and Suppl. Fig. 2).

**Figure 2.**
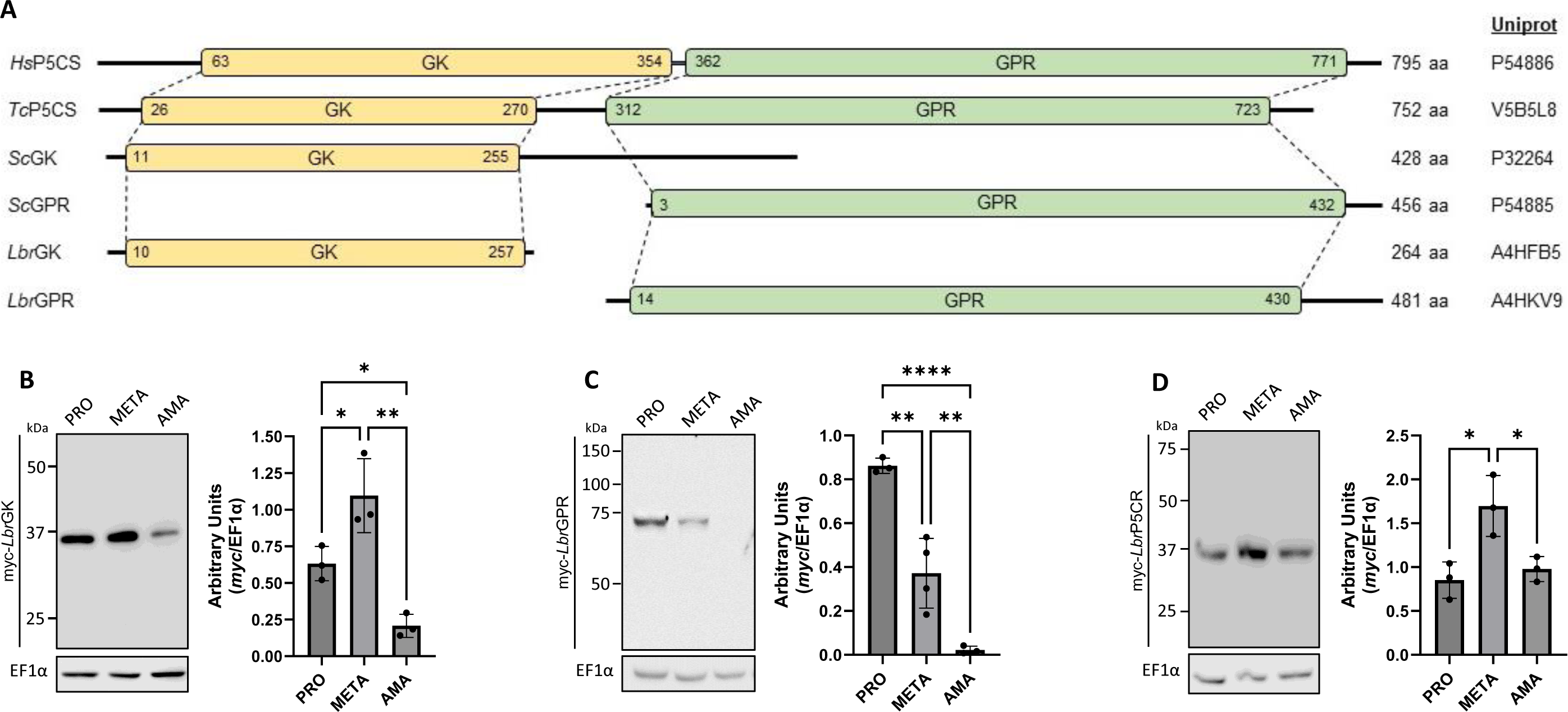
Expression of enzymes of the anabolic steps of the Glu-Pro pathway in different biological forms of *L. braziliensis*. A, Schematic illustration of the predicted locations of GK (yellow boxes) and GPR (green boxes) domains in *Homo sapiens* and *T. cruzi* P5CS, and in *Saccharomyces cerevisiae* and *L. braziliensis* GK and GPR enzymes. Numbers inside the boxes depict the residues where the domains start and end, according to InterPro(80). Codes shown on the right represent the UniProt IDs for each protein sequence shown. A full alignment is shown in Supplementary Figure 2. B, C and D, Expression of myc-*Lbr*GK (B), myc-*Lbr*GPR (C) and myc-*Lbr*P5CR (D) in procyclic (PRO) and metacyclic (META) promastigote forms, and in axenic amastigotes (AMA). Quantification of expression was done by dividing the intensity of the myc band signal by the signal of the loading control EF1α in non-saturated images. *P<0.05; **P<0.01; ****P<0.0001.

Given the ability of *L. braziliensis* promastigotes to recover their energy metabolism using only Glu or Gln and the fact that Pro is not essential for promastigote growth *in vitro* (20), it was likely that enzymes belonging to the Pro biosynthetic pathway are expressed in this life cycle stage. However, the expression of these enzymes in the infective metacyclic promastigote and amastigote stages is unknown, particularly given the preference of the amastigote stage to use glucose and fatty acids for energy production (23, 26, 27).

To assess the expression of GK, GPR and P5CR during the life cycle of *L. braziliensis*, we inserted N-terminal myc tags in each one of these genes and quantified the expression of these enzymes in procyclic and metacyclic promastigotes, as well as in axenic amastigotes (Figure 2). Our results show that GK and GPR, the enzymes responsible for catalysing the generation of P5C, present similar patterns of expression, being more highly expressed in promastigote forms, with higher expression in metacyclic and procyclic stages, respectively, but being largely absent in amastigotes (Figure 2A and B and Suppl. Fig. 3). P5CR, that catalyses the reduction of P5C into Pro, on the other hand, was more expressed in metacyclic cells in comparison to both procyclics and amastigotes, although its expression could readily be detected in all three biological forms. Moreover, P5CR was expressed in amastigotes at the same levels as in procyclic forms (Figure 2C and Suppl. Fig. 3). Additionally, mRNA for GK, GPR and P5CR was detected in procyclics, metacyclics and amastigotes at various levels, but were particularly upregulated in metacyclic forms (Suppl. Fig 4).

To verify the importance of the Pro anabolic steps of the Glu-Pro pathway, we created knockout (KO) cell lines for GK, GPR and P5CR and tested their ability to recover from nutritional stress utilising a single amino acid as Carbon source. First, interrupting the conversion of Glu into P5C by the individual knockout of GK and GPR did not impact the ability of *L. braziliensis* to use Glu or Gln as energy sources (Figure 3).

**Figure 3.**
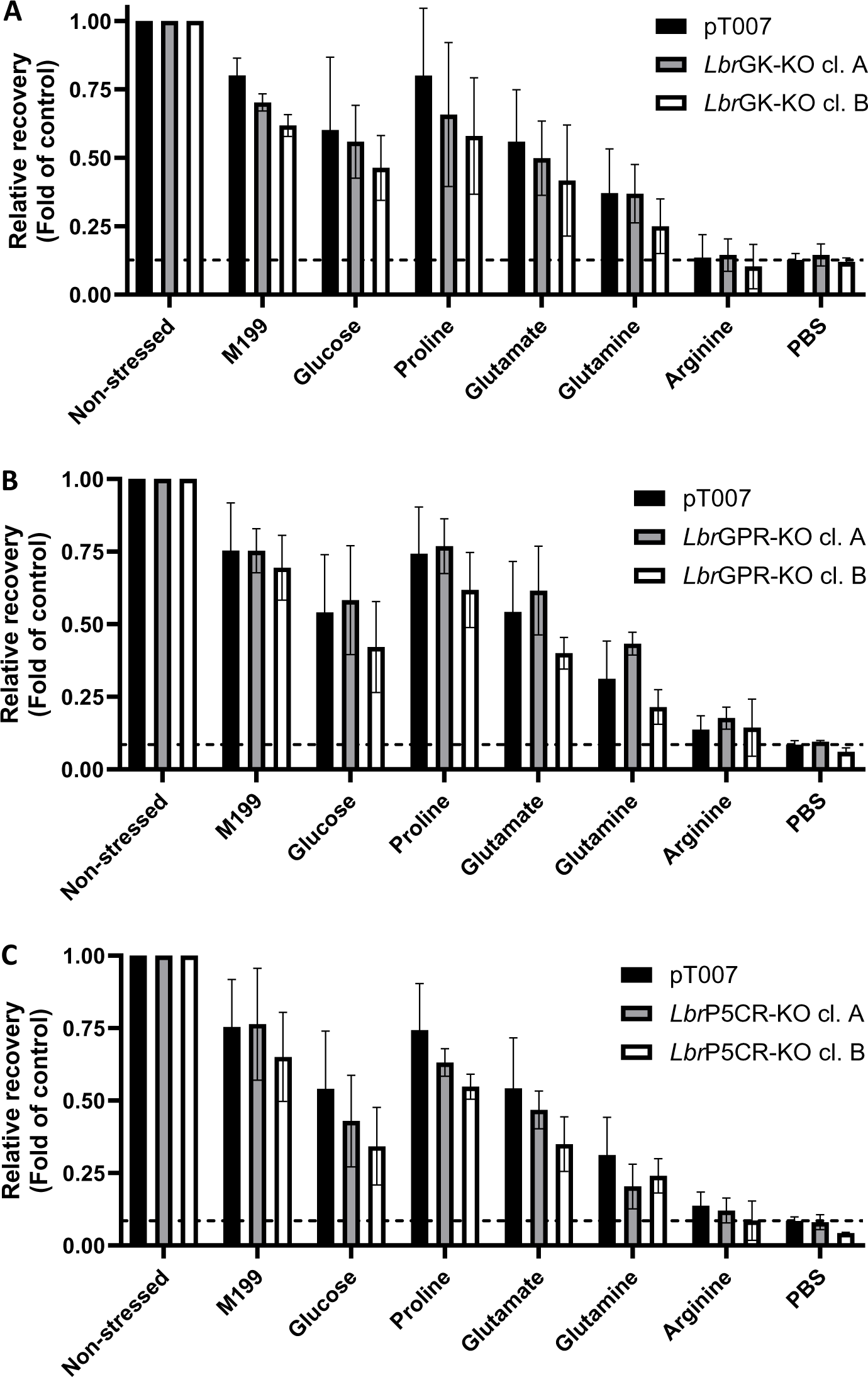
Recovery of cellular redox metabolism in cells knocked out for the anabolic enzymes of the Glu-Pro pathway subjected to nutritional stress. KO cell lines for *Lbr*GK (A), *Lbr*GPR (B) and *Lbr*P5CR (C) were subjected to nutritional stress in PBS and let to recover in nutritious buffer (M199 or PBS containing a single Carbon source). The level of MTT conversion was used as a measure for the recovery of cellular redox metabolism. The ratio of recovery in cellular metabolism was calculated in relation to the non-stressed group for each cell line, and the KO lines were compared to parental pT007 cells subjected to the same conditions by Two-way ANOVA.

In a myriad of organisms, the metabolism of Glu and Pro connect to the urea cycle by the conversion of P5C to ornithine by ornithine aminotransferase (OAT). Importantly, although urea has been identified as an excretion product for Leishmaniinae species (36, 37), it is most likely a product of the reaction catalysed by arginase to convert arginine into ornithine rather than an evidence of a functional urea cycle (4, 17). Furthermore, there is no evidence for OAT activity in trypanosomatids (4, 17). Nevertheless, we thus hypothesised that P5C could have been generated from Glu by a different, unconventional route and converted into Pro, which could explain the ability of the *Lbr*GK-KO and *Lbr*GPR-KO lines to recover cell redox metabolism using Glu and Gln. To test this possibility, we then knocked out P5CR and, to our surprise, verified that *Lbr*P5CR-KO cells are also able to efficiently utilise Glu and Gln to restore their redox metabolism (Figure 3C). Together, these data indicate that, different from what has been proposed for *T. cruzi* (10, 16), the Pro anabolic steps of the Glu-Pro pathway are not necessary for the obtention of energy from Glu or Gln in *Leishmania*.

### *L. braziliensis* does not utilize Glu to restore the intracellular pool of Pro

Phylogenetic analyses have shown a dynamic rate of lateral gene transfer from prokaryotes and fungi to *Leishmania* (38, 39) and, in *Leishmania*, such laterally transferred genes can be modified and undergo pseudogenization, degradation or loss (38). Genes laterally transferred to protists seem to be mostly involved in the metabolism of amino acids and carbohydrates (39). Importantly, many prokaryotes express an ornithine cyclodeaminase (OCD), which catalyses the conversion of ornithine into Pro and ammonia (40).

*Angomonas deanei* is the only trypanosomatid that expresses OCD, whose encoding gene was transferred to the protozoa from its obligatory endosymbiotic Betaproteobacteria (41, 42). Our searches in the Tritryp genome database did not retrieve any candidate gene for OCD in *Leishmania* spp. or *Trypanosoma* spp.. Moreover, to the best of our knowledge, the intermediate pyrroline-2-carboxylate generated by OCD has never been detected in metabolomic studies performed in various life cycle stages of either *Leishmania* or *Trypanosoma*. Yet, we raised the vague hypothesis that perhaps a highly modified OCD might have been incorporated by *L. braziliensis* during evolution and may be able to minimally replenish Pro levels under acute nutritional stress. To completely rule out this possibility, we subjected wildtype *L. braziliensis* promastigotes to nutritional stress in PBS for 2h and measured their ability to restore intracellular Pro levels using either Pro, Glu or Gln.

We observed a severe decrease (>80%) in the amount of intracellular free Pro in cells kept in PBS for 2h relative to cells collected prior to nutritional stress (Figure 4A), whereas cells maintained in M199 kept steady levels of intracellular Pro. At this time point, parasites were collected by centrifugation, washed once with PBS and resuspended in appropriate buffer containing either Pro, Glu or Gln for further 4h, after which the recovery of intracellular Pro levels were measured. A group of non-stressed cells followed the same protocol, but were always kept in culture medium. Importantly, such decrease in the level of free Pro should abolish the allosteric regulation Pro imposes on its biosynthetic pathway (25).

**Figure 4.**
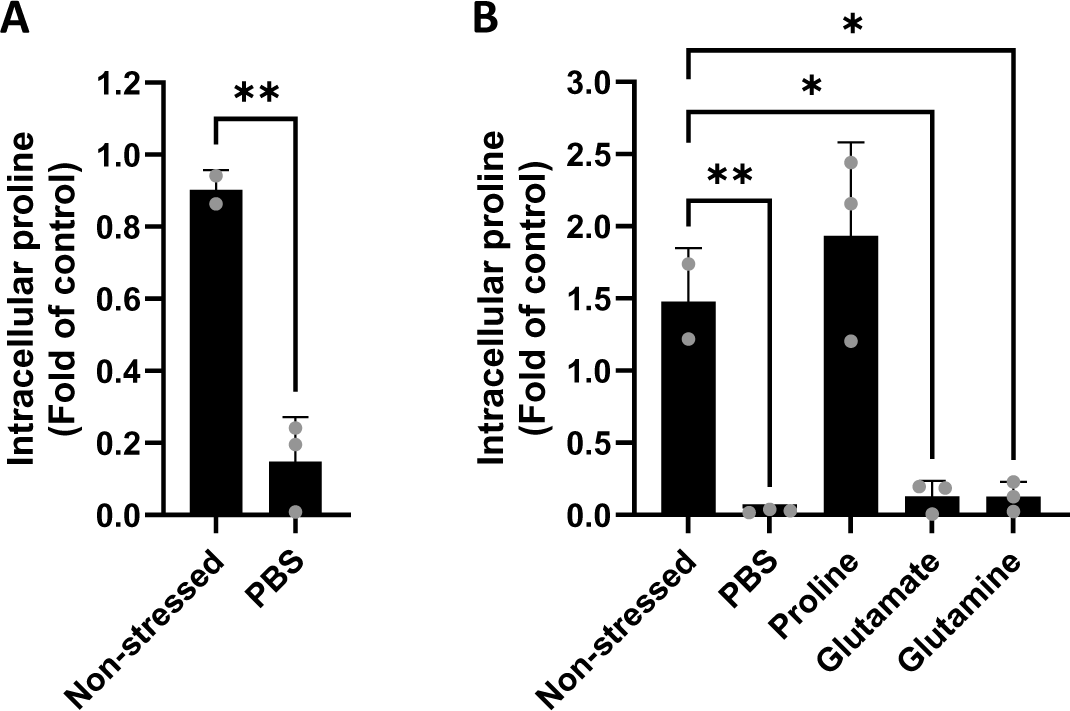
*L. braziliensis* wildtype cells do not use Glu and Gln to restore the intracellular pool of free Pro. (A) Wildtype *L. braziliensis* cells subjected to nutritional stress for two hours display a strong reduction in the amount of intracellular free Pro. (B) Cells let to recover for two hours reestablished the levels of intracellular free Pro when kept in presence of Pro, but not when in presence of Glu or Gln. The quantity of intracellular free Pro was calculated in comparison to cells taken prior to nutritional stress. Statistical analysis was performed by Student’s t-test (in A) or One-way ANOVA (in B) against the non-stressed control. *P<0.05; **P<0.01.

Interestingly, only cells that were let to recover from the nutritional stress in buffer containing Pro were able to restore their intracellular free Pro levels, whereas cells that were incubated in buffer containing either Glu or Gln were not able to do so and their level of free intracellular Pro was not significantly different from cells kept in PBS, but significantly reduced in comparison to non-stressed cells or cells recovered in Pro-containing buffer (Figure 4B). Noteworthy, cells that were allowed to recover in buffer containing Pro showed a tendence to have higher amounts of intracellular Pro (although this was not statistically significant), likely due to an increase in uptake stimulated by the nutritional stress (43–45). Remarkably, our results closely mirrored those obtained by Marchese (2017) in *L. amazonensis*, indicating that the biosynthesis of Pro from both Glu and Gln is not functional across *Leishmania* species.

### The knockout of key steps in the catabolic part of the Glu-Pro pathway reveal that Glu is an energy source even in the absence of Pro synthesis

We further considered the possibility that if generated at very low levels, Pro would be largely degraded by ProDH before accumulating in the cells. To address this possibility, we generated cells KO for *Lbr*ProDH and subjected them to nutritional stress and recovery using a single Carbon source. Interestingly, ProDH-KO cells were unable to recover their mitochondrial metabolism using Pro, as expected, but recovered their mitochondrial reductive metabolism using Glu or Gln likewise the parental cell line (Figure 5A). These results confirm that the recovery of mitochondrial metabolism using Glu and Gln does not require the biosynthesis of Pro in *L. braziliensis*.

**Figure 5.**
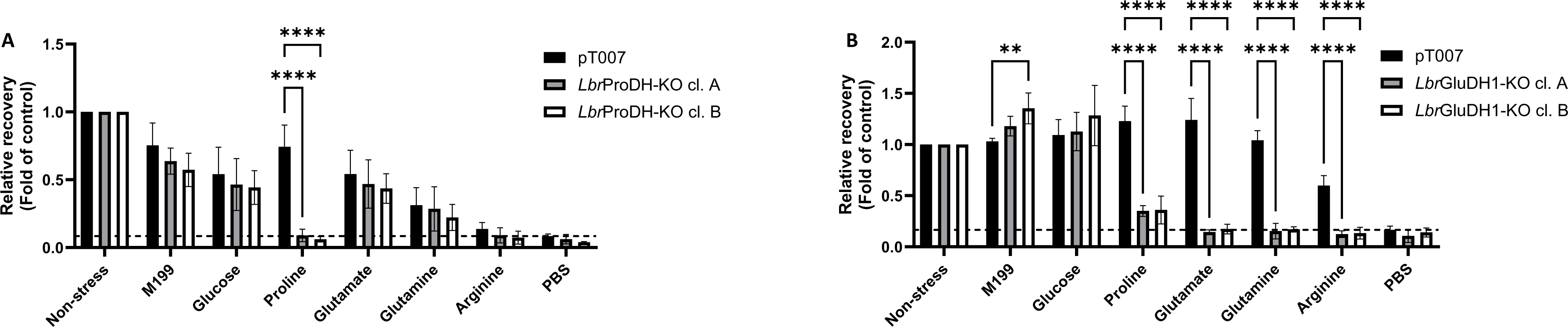
Recovery of cellular redox metabolism in cells knocked out for *Lbr*ProDH and *Lbr*GluDH1. KO cell lines for *Lbr*ProDH (A) and *Lbr*GluDH1 (B) were subjected to nutritional stress in PBS for four (*Lbr*ProDH-KO, A) or one (*Lbr*GluDH1-KO, B) hours and let to recover in nutritious buffer (M199 or PBS containing a single Carbon source). The level of MTT conversion was used as a measure of cellular redox metabolism.. The recovery in cellular metabolism was calculated in relation to the non-stressed group, and the KO lines were compared to parental pT007 cells subjected to the same conditions by Two-way ANOVA. **P<0.01; ****P<0.0001.

Given that Gln and Glu were efficiently utilised by the cell, we decided to investigate how these molecules were being catabolised. In most organisms, the main route for Glu catabolism is via GluDH. *Leishmania* parasites possess two genes annotated as GluDH, a longer version encoding for a protein of 1020 amino acids (here called *Lbr*GluDH1; LBRM2903_150017000) and a shorter version encoding for a protein of 452 amino acids (here referred to as *Lbr*GluDH2; LBRM2903_280038800). Of note, alignment of the *L. braziliensis* GluDH to the human and *S. cerevisiae* GluDH amino acid sequences showed that *Lbr*GluDH1 was most identical to *S. cerevisiae* GluDH2, whereas *Lbr*GluDH2 shows over 50% identity to *S. cerevisiae* GluDH1 and GluDH3 (Suppl. Table 2).

We generated cells knocked out for each version of GluDH and evaluated their recovery from nutritional stress using Pro, Glu and Gln as energy sources. First of all, we observed that GluDH1-KO cells were more sensitive to nutritional stress: after 4h in PBS, only a small proportion of cells was able to recover their metabolism, implicating in a reduction of about 75% in the conversion of MTT, when recovered in M199, and recovery using a single Carbon source was almost null in comparison to the parental line (Suppl. Figure 5B). Cells KO for GluDH2, on the other hand, did not have their recovery impaired and behaved likewise the parental line (Suppl. Figure 5C).

When GluDH1-KO cells were subjected to nutritional stress for 1h in PBS, they recovered in M199 in the same fashion as the parental line. In fact, at this time point, one clone displayed higher reductive potential than the parental line when recovered in M199; the reason for this phenomenon is not clear at this moment. Furthermore, we observed that GluDH1-KO clones were unable to restore their mitochondrial metabolism using Pro, Glu or Gln (Figure 5B). Intriguingly, when GluDH1-KO cells were subjected to nutritional stress for 2h they were able to recover in M199 likewise the parental line, but unable to efficiently use glucose or amino acids as energy sources (Suppl. Figure 5A). Given availability of ATP is required for the first steps of glycolysis, it is possible to hypothesise that GluDH1-KO cells survive on a more limitrophe metabolism and that their ATP levels fall below survival levels faster than in the parental line, which thus impairs the glycolytic metabolic flux after 2h of stress. Yet, recovery in a nutritionally diverse environment such as M199 is still possible.

Together, these results strengthen our understanding that the utilisation of Glu as energy source does not rely on its conversion into Pro prior to degradation, as *L. braziliensis* cells knocked out for ProDH could restore their reductive potential using Glu, but not using Pro. In turn, GluDH1-KO cells were unable to utilise both Glu and Pro as energy sources, evincing that the catabolism of both these amino acids rely on the activity of GluDH1, which further indicates that the catabolism of Pro follows a canonical route of degradation by the sequential activity of ProDH, P5CDH and GluDH.

In animals, GluDH predominantly localises in the mitochondrial matrix (46, 47), which indicates that the recovery of cellular redox capacity mediated by Glu during recovery from nutritional stress would require Glu to enter the mitochondrion prior to degradation. It has been shown, however, that the three forms of *Plasmodium falciparum* GluDH display distinct localisations, at least in intraerythrocytic stages: *Pf*GDH1 and 3 localise to the cytosol, while *Pf*GDH2 is present in the apicoplast (48, 49). Given that *Lbr*GluDH1 shows 30% of identity with *Pf*GluDH3, which is close to the 34.2% identity when compared to *Sc*GDH2 (Supplementary Table 2), we decided to add a C-terminal myc tag to *Lbr*GluDH1 and determine its intracellular localisation. We observed that the signal for *Lbr*GluDH1-myc superposes with the signal for the mitochondrial dye MitoTracker Deep Red (MTDR) (Supplementary Fig. 6), demonstrating its mitochondrial localisation.

Taking that *Lbr*GluDH1 is essential for the recovery of *L. braziliensis* from nutritional stress using Glu as sole Carbon source, its mitochondrial localisation implies that Glu enters the mitochondrion and is catabolised within the organelle, thus fuelling mitochondrial energy production. To test if Glu is capable of increasing mitochondrial activity, we exposed pT007 and *Lbr*ProDH-KO cl. A (in which the catabolism of Pro is interrupted) cells to nutritional stress for two hours followed by recovery in M199, PBS, PBS + Pro and PBS + Glu for two hours, and measured the accumulation of MTDR in each of these conditions by flow-cytometry. Of note, the accumulation of MTDR in actively respirating mitochondria has been previously reported (33, 34). A typical gating strategy is shown in Supplementary Figure 7A.

First, we exposed living and formaldehyde-fixed pT007 and *Lbr*ProDH-KO cells to MTDR for 30 minutes and verified that the living cells accumulated 3.5 to 4 times more MTDR than fixed cells on average (Supplementary Fig. 7B). Thus, we confirm that although MTDR stains the mitochondria in dead cells, its accumulation increases in actively respirating mitochondria.

Initially, we noticed that acute nutritional stress causes increased MTDR accumulation in pT007 cells: within two hours of incubation in PBS, cells accumulated approximately two times more MTDR than cells in the non-stressed group, which remained unchanged until the final time point, adding up to four hours of starvation (Figure 7A, B and Supplementary Fig. 7C). We reason this is likely due to mitochondrial swelling (50) rather than an increase in ΔΨ_m_ (51) upon starvation. Curiously, we did not observe this phenomenon in *Lbr*ProDH-KO cells (Figure 7C, D and Supplementary Fig. 7D).

After two hours of stress in PBS, cells were allowed to recover in nutritious environments. We noticed that cells transferred back into M199 accumulated MTDR to the same amounts as cells in the non-stressed group (Figure 6A – D). pT007 cells that were transferred to PBS + Pro, presented the highest fluorescence, whereas *Lbr*ProDH-KO incubated in PBS + Pro did not accumulate more MTDR than cells kept in PBS (Figure 6A – D). This confirms that Pro as highly capable of fuelling mitochondrial metabolism in *Leishmania*.

**Figure 6.**
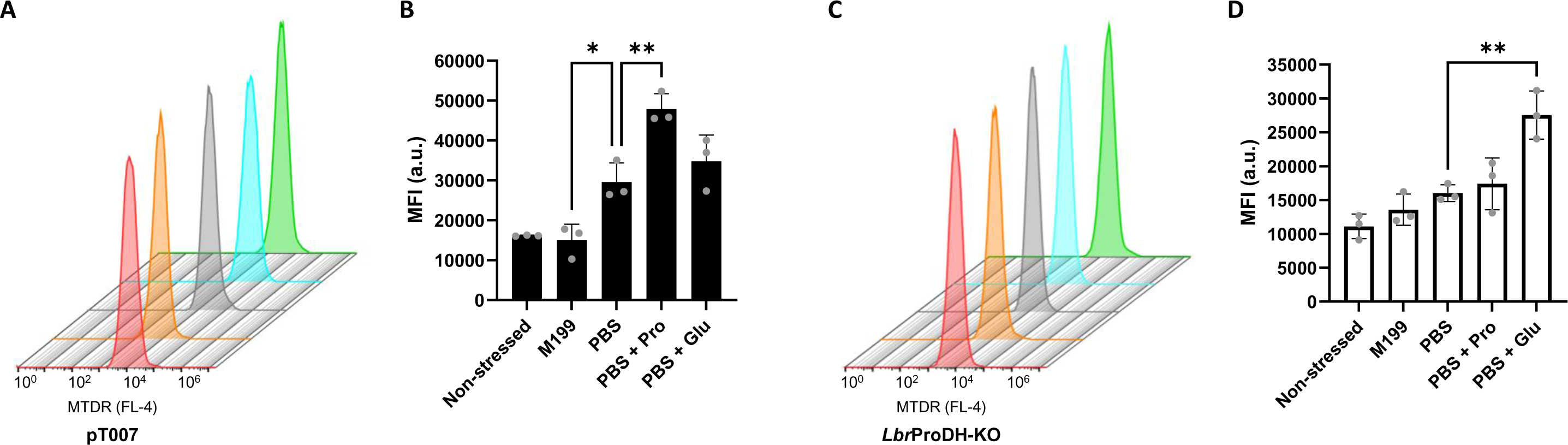
Catabolism of Proline and Glutamate increase mitochondrial membrane potential in *L. braziliensis*. (A, B) pT007 cells subjected to nutritional stress for two hours and allowed to recover in nutritious buffer accumulated MTDR at different levels. (C, D) *Lbr*ProDH-KO cells subjected to nutritional stress displayed a milder increase in MTDR accumulation under stress, and its recovery in buffer containing 3mM Glu caused a significant elevation in fluorescence. Quantification of fluorescence of *Lbr*ProDH-KO cells under different recovery conditions is shown in D. Typical histograms are shown in A and C for pT007 and *Lbr*ProDH-KO, respectively, and the quantification of fluorescence across three biological replicates are displayed in B and D. Red – non-stressed; Orange – starved and recovered in M199; Grey – starved and recovered in PBS; Blue – starved and recovered in PBS + Pro; Green – starved and recovered in PBS + Glu. *P<0.05; **P<0.01 by One-way ANOVA.

Interestingly, when pT007 cells were transferred to PBS containing Glu as single Carbon source, the accumulation of MTDR was higher than in cells transferred into M199, but not significantly different from cells maintained in PBS (Figure 6A, B). This phenomenon is most likely due to the proposed mitochondrial swelling that happens in cells under nutritional stress. *Lbr*ProDH-KO cells did not display increased MTDR accumulation during acute stress, on the other hand, they accumulated the highest levels of MTDR when incubated with 3mM of Glu (Figure 6C, D). This indicates an increase in intramitochondrial proton flux mediated by the catabolism of Glu.

Furthermore, the addition of FCCP during recovery from nutritional stress reduced the accumulation of MTDR in both pT007 and *Lbr*ProDH-KO cell lines in all conditions, including in the non-stressed group (Supplementary Figure 7E – N). This (1) further corroborates our observation that MTDR accumulation depends on ΔΨ_m_ and (2) confirms that the direct catabolism of Glu increases *Leishmania* ΔΨ_m_.

Altogether, these results show that the activity of *Lbr*ProDH is necessary for energy obtention from Pro, but not from Glu, whereas the activity of *Lbr*GluDH1 is essential for the efficient catabolism of both Pro and Glu. The recovery of energetic metabolism using these amino acids is due to elevation in electron transport chain activity with consequent increase in ΔΨ_m_, and predictable increase in ATP production.

### *Lbr*GluDH1-KO cells display higher amounts of intracellular free Glu under normal conditions, which is restored by Pro, but not by Glu, after nutritional stress

Given the inability of *Lbr*GluDH1-KO cells to restore their redox metabolism using either Pro, Glu or Gln, we sought to investigate how the intracellular pool of Glu was modulated under conditions of nutritional stress (1h) and recovery using a single Carbon source. *Lbr*GluDH1-KO cl. A was selected for these experiments.

First, we noticed that *Lbr*GluDH1-KO cells possessed 1.7 times higher amounts of intracellular Glu than the parental line under standard culture conditions (4.94 ± 0.45 and 8.47 ± 0.81 pM per cell in pT007 and *Lbr*GluDH1-KO, respectively) (Figure 7A). The level of intracellular Glu in both pT007 and GluDH1-KO lines decreased substantially (>60%) when these were incubated in PBS for one hour in comparison to cells that were resuspended and maintained in M199 for the same amount of time (Fig. 7B). At this point, the decrease in Glu levels was comparable between the two cell lines and reached a similarly low intracellular concentration (Suppl. Fig. 8A and B).

**Figure 7.**
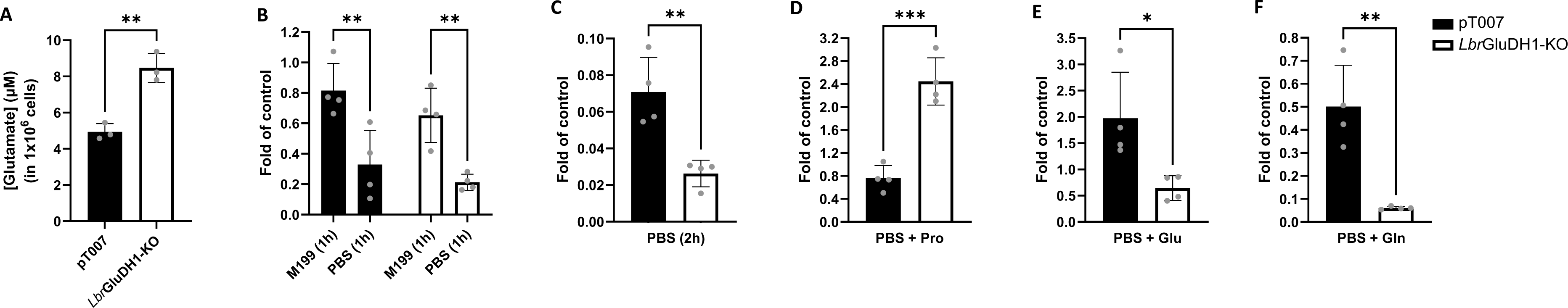
The role of *Lbr*GluDH1 in the recovery of intracellular free Glu after nutritional stress. (A) *Lbr*GluDH1-KO (white bars) cells display higher levels of intracellular Glu than the parental pT007 (black bars) cells under standard culture conditions. (B) The level of intracellular Glu decreases substantially in both pT007 and GluDH1-KO lines after one hour of stress in PBS in comparison to cells maintained in FBS-free M199. (C) After two hours of stress in PBS, the decrease in intracellular free Glu was more pronounced in *Lbr*GluDH1-KO cells. (D – F) After one hour of nutritional stress, both pT007 and *Lbr*GluDH1-KO were transferred to nutritious buffer for one hour: *Lbr*GluDH1-KO cells recovered in buffer containing 3mM Pro accumulated more intracellular Glu than the pT007 line (D), when incubated in buffer containing 3mM Glu, pT007 cells accumulated Glu beyond what was seen in cells under normal conditions, whereas the *Lbr*GluDH1-KO was not capable of restoring its pool of intracellular free Glu (E). (F) Incubation of pT007 cells with 3mM Gln after nutritional stress did not restore the levels of Glu, and this was further reduced in *Lbr*GluDH1-KO cells. *P<0.05; **P<0.01; ***P<0.001.

After stress for one hour, cells were washed once in PBS and then transferred to PBS only or PBS containing either Pro, Glu or Gln as a single Carbon source for another hour, after which the intracellular levels of Glu were measured. As expected, further incubation in PBS caused a further decrease in intracellular levels of Glu in both cell lines, and even though this effect was more pronounced in *Lbr*GluDH1-KO cells (Fig. 7C and Suppl. Fig. 8D, H), reasonably due to the higher Glu levels this cell line displays under normal conditions, the concentration of intracellular Glu was similar in both cell lines (Suppl. Fig. 8C). When transferred to PBS containing Pro, both cell lines were able to at least partially restore their pool of intracellular Glu: intracellular Glu increased by 2.31-fold in pT007 (in comparison to cells stressed in PBS for one hour), which elevated it to almost the same level as measured in cells under normal conditions, and by 11.52-fold in *Lbr*GluDH1-KO cells, leading to an accumulation of this amino acid, which was 2.44 times higher than that observed for GluDH1-KO in standard conditions (Fig. 7D and Suppl. Fig. 8E, I). This corroborates the idea that the catabolism of Pro follows a canonical route in *Leishmania*: in the absence of *Lbr*GluDH1, the Glu generated by the activity of ProDH and P5CDH accumulates within the cell.

As expected, increases in intracellular levels of Glu were observed for both cell lines when transferred from PBS (1h) to PBS containing Glu (5.99- and 3.03-fold in pT007 and *Lbr*GluDH1-KO, respectively) (Suppl. Fig. 8F, J). Strikingly though, while the pT007 cells accumulated Glu at 1.97-fold higher than the cells collected at the beginning of the experiment, most likely due to increased Glu uptake, the level of Glu in *Lbr*GluDH1-KO cells never reached that observed for cells under steady state (0.64-fold) (Figure 7E), indicating that the absence of *Lbr*GluDH1 causes a delayed response to Glu starvation, and a consequent deficient upregulation of Glu transporting activities.

Furthermore, we observed that incubation of pT007 cells with Gln after nutritional stress did not restore the levels of Glu to the levels observed prior to incubation in PBS, but rather just precluded a further decrease in Glu levels (Fig. 7F and Suppl. Fig. 8G). Several potential explanations could be raised for this phenomenon, such as (1) slow level of Gln uptake, (2) rapid consumption/incorporation of Gln by other metabolic routes leaving only a small amount of free Gln that could be converted into Glu, and/or (3) slow rates of conversion of Gln into Glu (likely via glutaminase) (52). Neither of these, however, seem to be a likely explanation for the inefficiency of *Lbr*GluDH1-KO in replenishing (or at least avoid further decrease in) intracellular Glu levels: in this cell line, a decrease of 3.51-fold in Glu concentration was observed in cells incubated in PBS + Gln in comparison to cells starved in PBS for 1h (Fig. 7F and Suppl. Fig. 8K). Yet, the level of Glu in *Lbr*GluDH1-KO cells incubated in PBS + Gln was not as low as that observed in *Lbr*GluDH1-KO cells kept in PBS for two hours (Suppl. Fig. 8L). These results show a deficiency of GluDH1-KO cells to efficiently obtain/utilise Gln.

Taken together, these results show that *Lbr*GluDH1 is the major route for Glu catabolism in *L. braziliensis* and also seem to suggest that the activity of *Lbr*GluDH1 is an important signal involved in triggering the expected increase in Glu transport in response to nutritional stress.

### *Lbr*P5CR may play moonlight functions

We observed that the anabolism of Pro from Glu is likely not functional in *Leishmania* spp. and neither of the proteins (*Lbr*GK, *Lbr*GPR and *Lbr*P5CR) involved in this pathway are essential *per se* for *in vitro* growth and recovery from nutritional stress (Figure 4). Notably, *Lbr*P5CR was expressed in both procyclic and amastigote forms at equivalent levels and its expression in amastigote forms was dissonant to the expression of its upstream enzymes *Lbr*GK and, most noticeably, *Lbr*GPR (Figure 3 and Suppl. Figure 1). Although the level of expression does not necessarily correlate with the level of activity of a given enzyme, such dissonance and the fact that many metabolic enzymes play moonlighting functions in eukaryotic cells, particularly regarding RNA binding activities (53–55), prompted us to investigate potential divergent functions played by *Lbr*P5CR in the promastigote and amastigote stages.

Importantly, enzymes that participate of the Glu-Pro pathway, namely ProDH and P5CDH, were identified in an mRNA-bound proteome of *L. mexicana* (56). A potential moonlight function as a ‘mRNA stabilizing’ protein has been reported for *T. brucei* P5CR in an artificial tethering assay (57). Thus, to check whether *Lbr*P5CR would interact with different groups of proteins in procyclic and axenic amastigote forms, myc-*Lbr*P5CR was immunoprecipitated from both developmental stages (Suppl. Figure 9) and subjected to mass spectrometry for the identification of its putative interaction partners.

A total of 334 and 277 potential interactors of myc-*Lbr*P5CR were identified in promastigotes and amastigotes, respectively: of these, 40 and 48 proteins were common between the three biological replicates in each biological form, respectively, and were considered for further analysis (Figure 8A, B). Interestingly, only four proteins were identified as putative myc-*Lbr*P5CR in both biological forms, whereas 36 proteins were exclusively identified in sample eluates from procyclic forms, and 44 in amastigote samples (Figure 8C).

**Figure 8.**
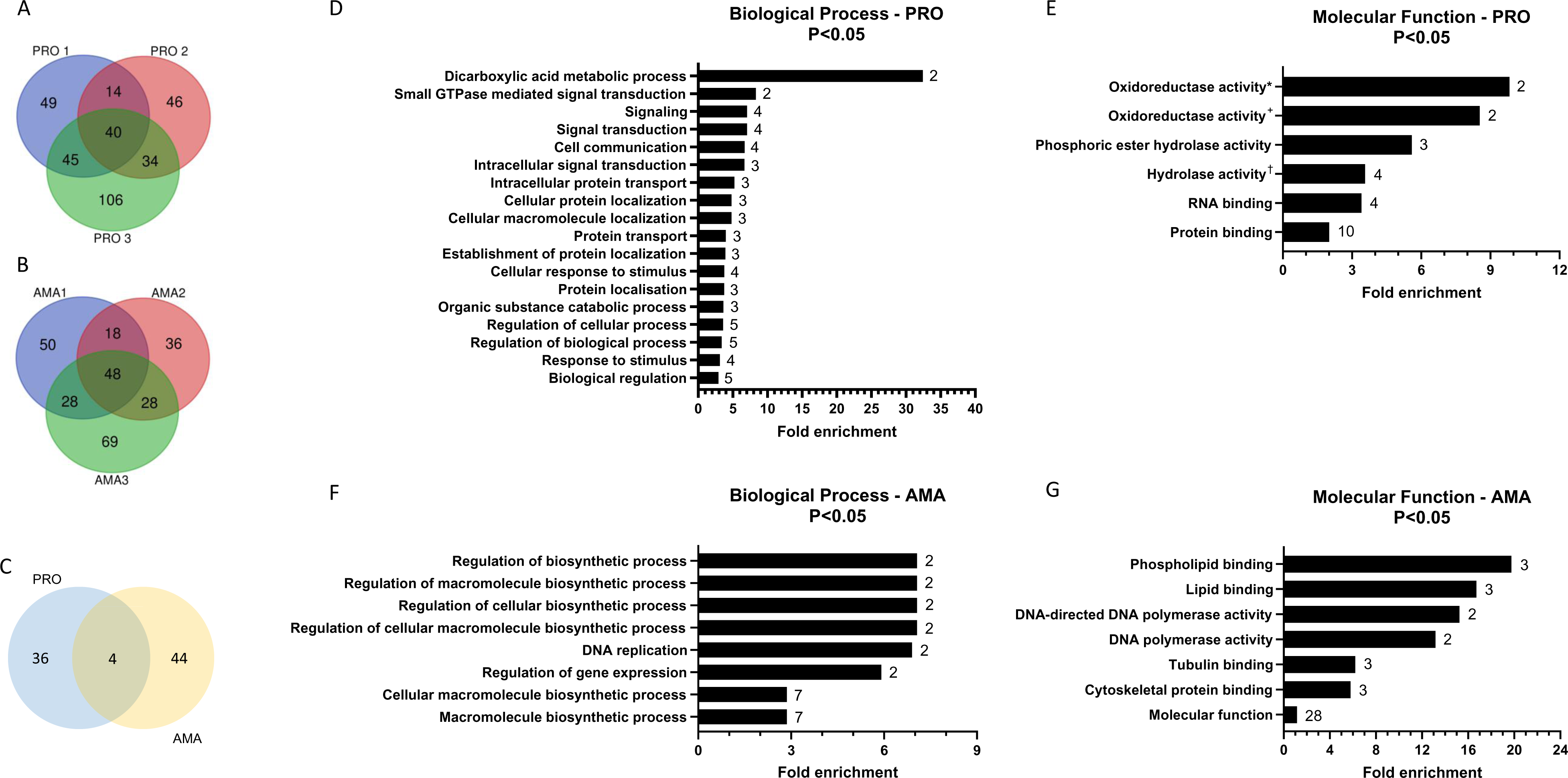
GO Term analysis of putative myc-*Lbr*P5CR partners of interaction identified by Mass Spectrometry. Proteins identified in the three biological replicates of myc-*Lbr*P5CR immunoprecipitation from procyclic (A) and amastigote (B) forms were compared. Proteins that were common among the three biological replicates were used to identify proteins specific to each stage: 36 proteins were identified only in in procyclic forms and 44 only in amastigotes (C). (D – G) These were subjected to GO Term analysis for Biological Process (D and F) and Molecular Function (E and G) in TritrypDB. Only GO Terms that displayed *p* < 0.05 and contained at least two genes were considered. Numbers in front of the bars reflect the number of genes identified in each GO Term.

The putative myc-*Lbr*P5CR interactors that were specific to each of the biological forms were subjected to gene ontology (GO) term analysis regarding Biological Process and Molecular Function in TriTrypDB (Figure 8D – G). Only GO Terms that contained two or more genes and with *p* values <0.05 were considered in the analysis. In procyclic forms, the GO Terms determined for myc-*Lbr*P5CR interaction partners are mostly related to metabolic processes, protein transport and localisation, and response to stimulus and signaling. Importantly, myc-*Lbr*P5CR from procyclic cells interacted with ProDH, malate dehydrogenase and glycerol-3-phosphate dehydrogenase, all of which are involved in energetic metabolism and maintenance of adequate redox balance. Interestingly, four proteins with a molecular function of “RNA binding” were identified among myc-*Lbr*P5CR interaction partners and a cross-analysis against orthologue RNA-binding proteins (RBPs) identified in the global *Leishmania mexicana* mRBPome (56) revealed other five RBPs that were mostly enriched in the mRBPome of procyclic promastigote forms. As such, proteins related to RNA binding activities account for a quarter (9 out of 36 proteins) of all putative myc-*Lbr*P5CR interaction partners in *L. braziliensis* procyclic forms (Table 1).

**Table 1.** RBPs identified as putative myc-*Lbr*P5CR interaction partners.

| Gene ID | Predicted function | RNA binding |  |
| --- | --- | --- | --- |
|  |  | GO Term | mRBPome* |
| LBRM2903_150019900 | Eukaryotic translation initiation factor 4 gamma type 2, putative | X |  |
| LBRM2903_150022000 | cAMP specific phosphodiesterase, putative |  | X |
| LBRM2903_170007000 | Tetratricopeptide repeat/TPR repeat, putative |  | X |
| LBRM2903_190008700 | Fibrillarin, putative | X | X |
| LBRM2903_260022000 | proline oxidase (ProDH), mitochondrial precursor-like protein |  | X |
| LBRM2903_280010000 | DIS3-like exonuclease, putative | X |  |
| LBRM2903_310015200 | Mitochondrial RNA binding complex 1 subunit, putative | X | X |
| LBRM2903_310015300 | hypothetical protein |  | X |
| LBRM2903_310015400 | mitochondrial RNA binding protein, putative |  | X |

\*Proteins identified in the *L. mexicana* mRBPome (56) and whose enrichment was highest in procyclic forms (cross-linked or not cross-linked).

GO Terms that contained at least two putative myc-*Lbr*P5CR interaction partners in amastigote forms were mostly related to regulation of biosynthetic processes and macromolecule (protein, lipid, and nucleic acids) binding. This includes two (one catalytic and one regulatory) subunits of DNA polymerase II, the RNA-polymerase associated CTR9 protein, one mitochondrial DEAD box-containing protein, a 4E-interacting protein, a asparaginyl-tRNA synthetase, and three ribosomal proteins, indicating that myc-*Lbr*P5CR from amastigote forms might be somehow involved in nucleic acid metabolism and mRNA translation. Taken together our results indicate that *Lbr*P5CR may play moonlight functions in *L. braziliensis* and that these are divergent between the procyclic and amastigote stages.

## Discussion

Trypanosomatids diverged from other eukaryotes between 200 and 500 million years ago, a period marked by the emergence of Arthropoda and Mammalia and, with them, the appearance of new niches (58, 59). As trypanosomatids adapted to a parasitic lifestyle, events of genome streamlining with loss of many protein coding genes accompanied by the expansion of gene families related to parasitism and nutrient acquisition, particularly of purine and amino acid transporters, occurred (60). Moreover, as trypanosomatids adapted to a dixenous life cycle, novel mechanisms to respond and adapt to the different environments encountered within vertebrate and invertebrate hosts became necessary (61). As a result, different morphologically and metabolically adapted biological forms appeared. For example, once ingested by the phlebotomine vector, the slow growing, mammalian-infective, sugar- and fatty acid-consuming *Leishmania* amastigote forms, in which the functionality of the TCA cycle seems to be reduced and mostly redirected to provide intermediates for anabolic processes, switch their metabolism to become fast dividing and highly metabolic active promastigote forms, in which the consumption of glucose and amino acids are elevated and the TCA cycle is used for energy production (23, 27).

Moreover, trypanosomatids are able to sense and progressively regulate their metabolism to compensate for the lack of nutrients. When starved for amino acids, these parasites activate a metabolic response that upregulates the expression and the activity of amino acid transporters (43–45). Of these, proline transporters seem to be specially upregulated under amino acid starvation conditions (44), likely due to the high availability of this amino acid in haemolymph of the insect vector (62), which correlates with Pro being the preferred amino acid used for energy production under glucose-limited conditions (14–16). In *Leishmania*, Pro is internalised via three separate transporting systems: systems A, B and C, of which systems A and B are expressed in promastigote forms and system C is amastigote-specific. Of note, whereas system B seems to be more specific for Pro, alanine, and, likely, cysteine, system A (AAP24) is a broad specificity transporter and also responsible for the bulk of proline uptake in procyclic forms (63–65), whereas the transport of Glu and Gln is more limited: Glu transport is (mostly) mediated by a single system, which may also carry Gln (66), as in *T. cruzi* (67), although other transporters seem to carry Glu at low levels (68). The transport of Gln has never been directly characterised in *Leishmania*, and in *T. cruzi*, this was found to be mediated by a single transporting system as well (69).

Once inside the cell, these nutrients can be catabolised. Here, we show that the catabolism of glucose and Pro efficiently recovers *Leishmania* cellular metabolism after nutritional stress, as expected, and we further show that Glu and, to a lesser extent, Gln can also be utilised in this process. In *T. cruzi*, Glu is also employed as energy source, but not without being first converted into Pro (via the activity of P5CS and P5CR), which is imported into the mitochondrion and catabolised in a canonical pathway (4, 10, 16). *Leishmania* species retain all genes necessary for a functional Pro biosynthesis pathway (70) and at least one of them (GK) has been demonstrated to be functional in *Leishmania donovani* (25). Here, we showed that the three enzymes involved in the anabolism of Pro from Glu are expressed at similar levels in procyclic forms of *L. braziliensis*, and that their expression fluctuates across life cycle stages, which might be associated with different functions played by these proteins in each life stage. When procyclic forms were subjected to nutritional stress, a significant drop in the levels of intracellular free Pro was observed, and this was only replenished when cells were allowed to recover in proline-containing buffer. The fact that incubation with neither Glu nor Gln could restore intracellular Pro levels suggested that the anabolic part of the Glu-Pro pathway has limited functionality in *L. braziliensis*.

In fact, it has been shown that, in *Leishmania*, most of the internalised Glu is either retained as a free amino acid or incorporated into TCA cycle intermediates (23, 24). It could be hypothesised that the use of Glu by *Leishmania* parasites follows the same route as in *T. cruzi* and that the absence of isotope-labelled Pro in procyclic forms fed with ^13^C-U-Glu is because the catabolic part of the Glu-Pro pathway consumes all the Pro generated by the anabolic steps before it could be accumulated within the cell. We further addressed the possibility of accelerated consumption of Pro by generating *Lbr*ProDH-KO lines and observed that although these cells were no longer capable of utilising Pro as an energy source, the use of both Glu and Gln was unchanged, demonstrating that the biosynthesis of Pro is not a requirement to reestablish cell redox metabolism using either Glu or Gln. The inability of *L. braziliensis* to synthesise Pro from other amino acids is in agreement with the necessity of Pro supplementation to sustain *L. braziliensis* growth *in vitro* (19).

Assuming that the catabolic steps of the *Leishmania* Glu-Pro pathway work similarly to the that observed in trypanosomes, it seems clear why the catabolism of Pro recovers the redox metabolism more efficiently than the catabolism of Glu: as Pro enters the mitochondrion, it is catabolised to P5C by ProDH, with the reduction of FAD to FADH_2_, P5C is spontaneously converted into GSA, and GSA is oxidised into Glu by P5CDH alongside the reduction of NAD^+^ to NADH. Both FADH_2_ and NADH can directly feed the mitochondrial electron transport chain and fuel ATP production. Additionally, Glu will be catabolised into α-ketoglutarate and participate in the anaplerosis of the TCA cycle, further stimulating energy production (10, 15, 16, 71, 72). Future biochemical characterisation of each one of the enzymes involved in Pro catabolism is necessary to answer if this hypothesis holds true.

It is noteworthy saying that it is not clear, at this point, whether the production of FADH_2_ and NADH during Pro oxidation is the only factor that makes Pro a preferred energy source over Glu. It is likely that the high-capacity Pro transporters can rapidly replenish the starving cells with this amino acid, which facilitates its catabolism and makes it more efficient in recovering the *Leishmania* redox metabolism than Glu, whereas Glu uptake is either rather inefficient or is not as upregulated as Pro transporters under nutritional stress (44). The recent identification of genes encoding for putative Glu transporters (particularly LmjF.22.0230 and its orthologues) (68) shall help to elucidate this possibility.

Another step that can limit the efficiency of a given amino acid in restoring energy metabolism is its ability to enter the mitochondrion. In *T. cruzi*, the presence of *Tc*ProDH in the mitochondrial inner membrane implies the existence of a (still uncharacterised) transporting mechanism that takes this amino acid into the organelle (72). Here, we showed that *Lbr*GluDH1 is essential for the catabolism of both Pro and Glu, and that it is also a mitochondrial enzyme, as occurs in most organisms (7). As such, it is necessary for both these nutrients enter the mitochondrion before its catabolism. As for the moment, there is no evidence that Glu can be transported into the mitochondria of *Trypanosoma* spp., whereas our data demonstrate this happens in *Leishmania*. Determining whether Glu enters the mitochondrion via a mitochondrial Glu transporter, as has been shown in other organisms (5, 7, 73), or as part of the malate-aspartate shuttle will require future investigation. Nonetheless, these results highlight a significant difference in the metabolism of amino acids between *Trypanosoma* and *Leishmania* and suggests that a diversification in amino acid catabolic pathways may exist within Trypanosomatidae.

The inability of *Lbr*ProDH-KO cells to catabolise Pro indicated that the catabolic part of the Glu-Pro pathway was functional. To confirm this, we knocked out the final step of the pathway (*Lbr*GluDH1), and verified that, as expected, cells became unable to utilise Pro, Glu and Gln for energy production. Moreover, we observed that Pro could be used to replenish the pool of free Glu in *L. braziliensis* after nutritional stress, and that Glu generated from Pro accumulated inside the cell when *Lbr*GluDH1 was knocked out. These results confirmed that the catabolic steps of the Glu-Pro pathway are functional, and that the generation of Glu from Pro is possible, as had been suggested before (21, 37, 74, 75).

Strikingly though, we observed that *Lbr*GluDH1-KO cells were less capable of restoring the levels of intracellular Glu when supplied with Glu or Gln during recovery from nutritional stress than the parental pT007 line. Glu can be obtained by active transport from the buffer or by the catabolism of Gln (4, 66, 68, 76, 77). It has been shown that inhibition or knockdown of GluDH is associated with a decrease in the transport of both Glu and Gln (78, 79). Given the activity of *Leishmania* GluDH gradually increases towards the production of α-ketoglutarate under starvation conditions (37), signifying that GluDH is part of a metabolic response to starvation, it is possible, thus, that the activity of GluDH is necessary to trigger an appropriate response to starvation and stimulate Glu and Gln uptake.

Based on the results presented here, we propose that the Glu-Pro pathway in *Leishmania* differs from the well characterized pathway in *Trypanosoma* in several aspects (Figure 9): (1) *Leishmania* does not possess a bifunctional P5CS, but does possess genes encoding for GK and GPR; (2) The functionality of the anabolic steps of the pathway seems to be limited, if functional at all, and as such, (3) *Leishmania* does not require the biosynthesis of Pro from Glu to use the latter as an energy source. In fact, (4) Glu can likely be transported into the mitochondrion and directly catabolised as a means of increasing mitochondrial function and, therefore, propelling energy production.

**Figure 9.**
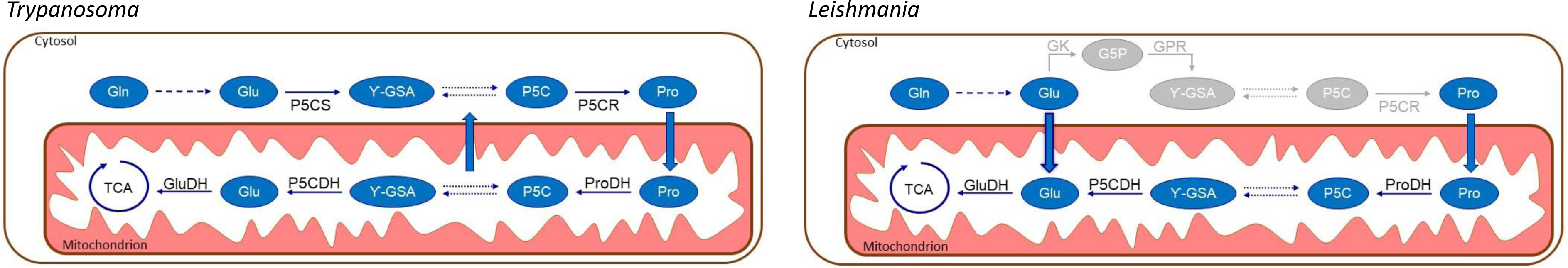
Comparison between the proposed models for the functionality of the Glu-Pro pathway in *Trypanosoma* and *Leishmania*. While trypanosomes possess a bifunctional P5CS enzyme, *Leishmania* displays genes encoding for GK and GPR. The activity of P5CS followed by the activity of P5CR can generate Pro from Glu in *Trypanosoma* spp., but this pathway seems to have limited functionality in *Leishmania* species. In both cases, Pro is primarily degraded by ProDH. The production of P5C from Pro and its degradation via P5CDH has been shown in trypansomes: knockdown of P5CDH in *Trypanosoma* leads to na accumulation of this metabolite in the mitochondria and its diffusion to the cytosol, but this has never been demonstrated in *Leishmania.* The activity of P5CDH generates Glu, which is catabolised into α-ketoglutarate, which enters the TCA cycle. We showed here that *Leishmania* the Glu serving as substrates for GluDH1 can be either generated from Pro or, most strikingly, (likely) directly imported into the mitochondrion via an unknown transporter. The catabolism of Glu can increase mitochondrial membrane potential.

## Acknowledgments

We thank Dr. Arthur A. C. de Oliveira (FFCLRP, USP) for providing phosphoenolpyruvic acid and Dr. Thiago A. Cunha (FMRP, USP) for donating polypropylene plates. We also thank Dr. Jennifer A. Black (FMRP, USP) for her assistance with FACS data analysis, and Msc. Tânia Paula A. Defina (FMRP, USP) for the RT-qPCR data. We thank Prof. Ariel M. Silber and Dr. Letícia Marchese (ICB, USP) for the exchange of thoughts and protocols. We also would like to thank the VEuPathDB team for maintaining incredible databases and tools.

