## Supplementary Figures for "Genetic and functional dissection of the glutamate-proline pathway reveals a shortcut for glutamate catabolism in *Leishmania*"

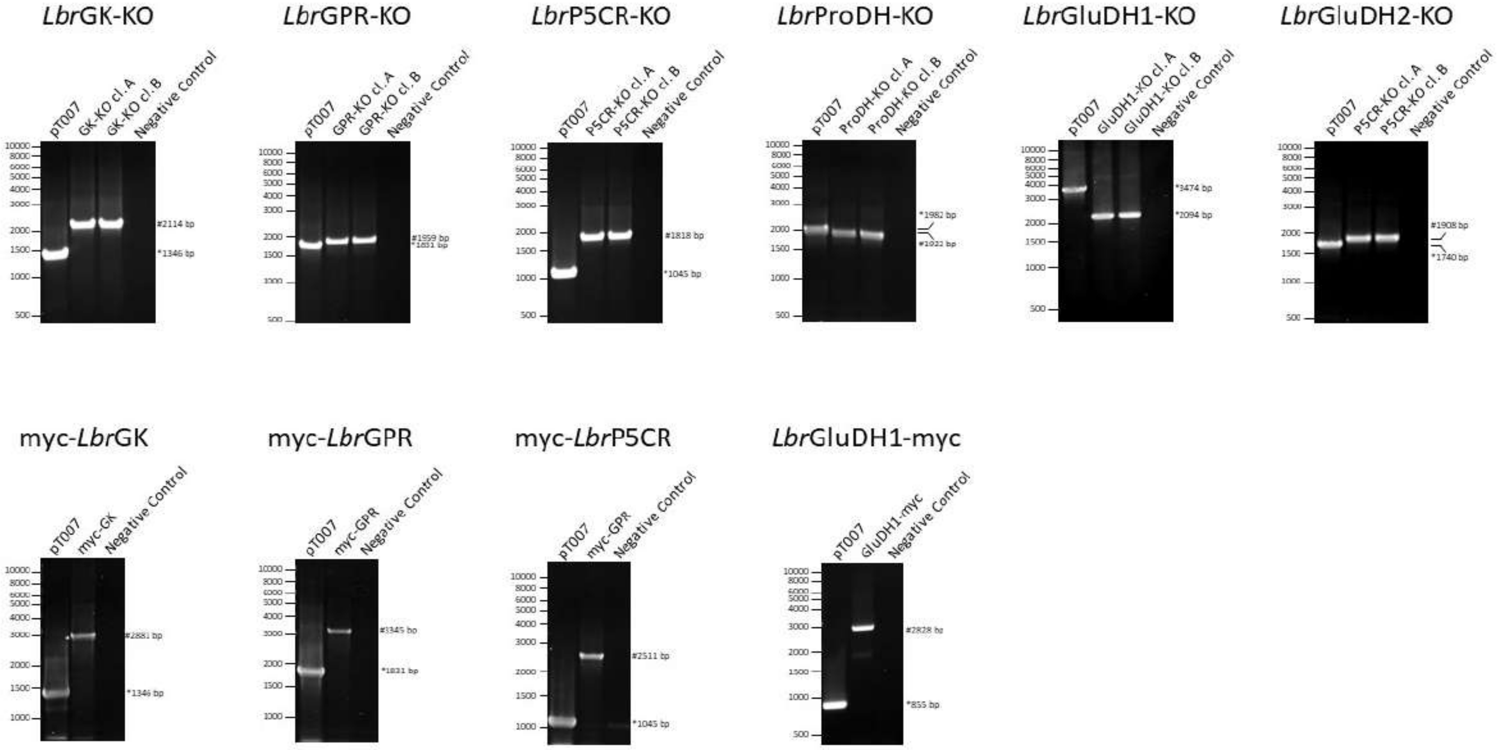


Supplementary Fig. 1

Supplementary Figure 1. Examples of PCRs to confirm knockout and tag of *Lbr*GK, *Lbr*GPR, *Lbr*P5CR, *Lbr*ProDH, *Lbr*GluDH1 and *Lbr*GluDH2. *Native alleles; #Modified alleles.

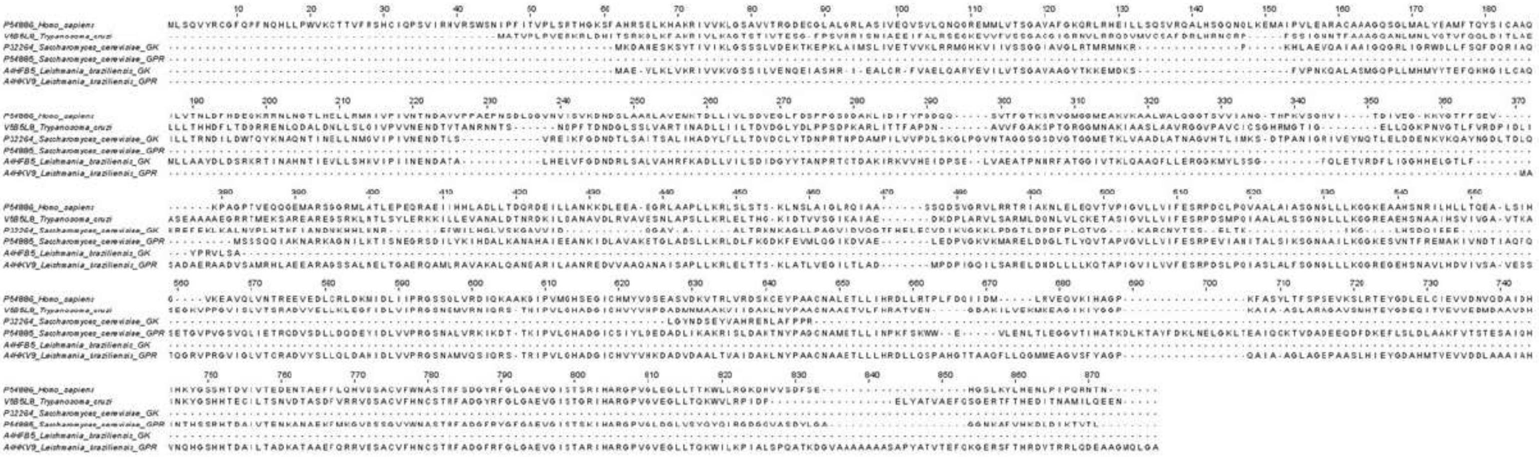

Supplementary Fig. 2

Supplementary Figure 2. Full alignment of GK, GPR and P5CS proteins from different species. Sequences were aligned with Clustal Omega.

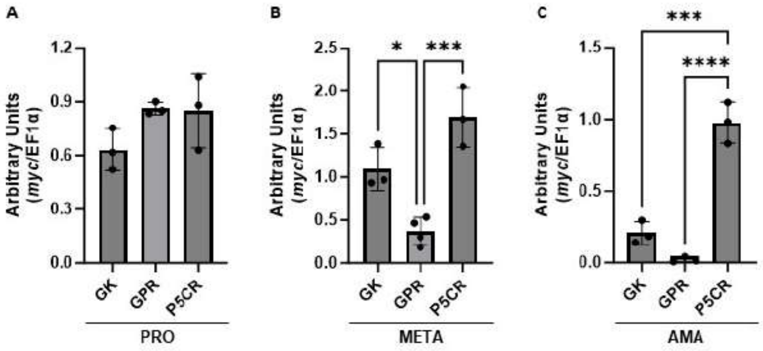


Supplementary Fig. 3

Supplementary Figure 3. Comparative analysis of the expression of myc-*Lbr*GK, myc-*Lbr*GPR and myc-*Lbr*P5CR in different biological forms of *L. braziliensis*. Data shown in Figure 3 were replotted to compare their expression in each procyclic (A), metacyclic (B) and axenic amastigote (C) forms individually. *P<0.05; ***P<0.001; ****P<0.0001.

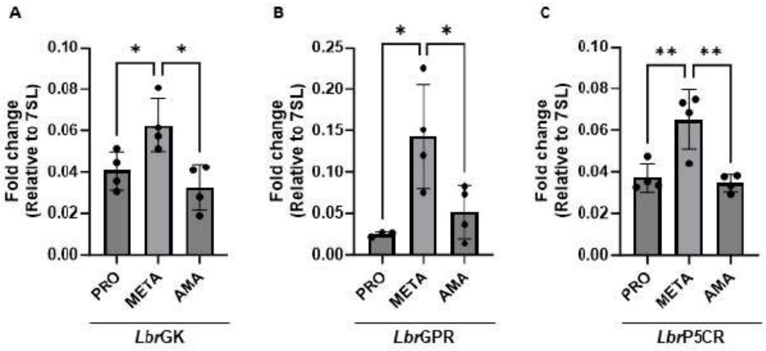


Supplementary Fig. 4

Supplementary Figure 4. Analysis of the level of mRNA encoding for *Lbr*GK, *Lbr*GPR and *Lbr*P5CR in different biological forms of *L. braziliensis.* Quantification of the transcripts encoding for *Lbr*GK (A), *Lbr*GPR (B) and *Lbr*P5CR (C) was performed by RT-qPCR in each biological form obtained from axenic culture. The splice leader (7SL) was used as a normaliser. *P<0.05; **P<0.01.

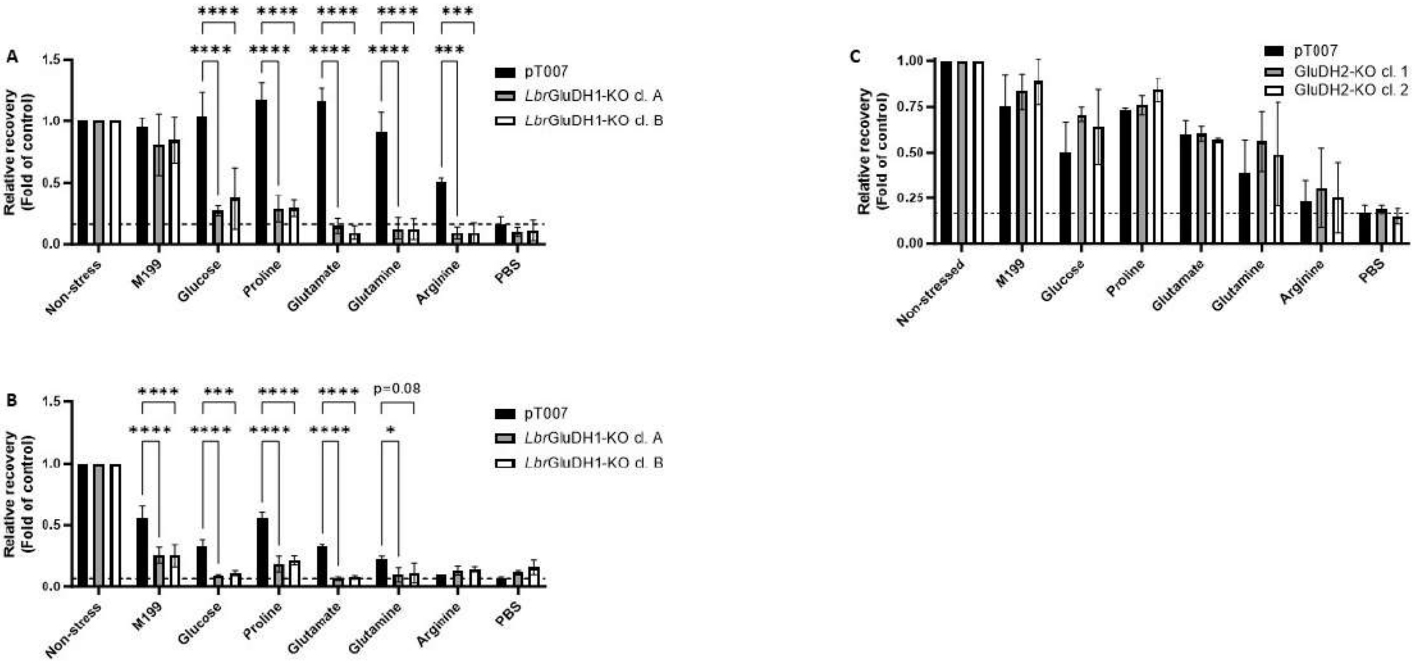


Supplementary Fig. 5

Supplementary Figure 5. *Lbr*GluDH1-KO cells are more sensitive to nutritional stress. *Lbr*GluDH1-KO cells subjected to nutritional stress for two (A, *n* = 3) or four (B, *n* = 2) hours in PBS displayed reduced capacity to reestablish their redox metabolism using a single Carbon source. When subjected to nutritional stress for two hours, *Lbr*GluDH1-KO lines were able to recover their cellular metabolism in the complex M199, but not in buffer containing a single Carbon source. (C) Cells KO for GluDH2 recovered from nutritional stress in the same fashion as the parental cell line (*n* = 2). The recovery in cellular metabolism was calculated in relation to the non-stressed group, and the KO lines were compared to parental pT007 cells subjected to the same conditions by Two-way ANOVA. *P<0.05; ***P<0.001; ****P<0.0001.

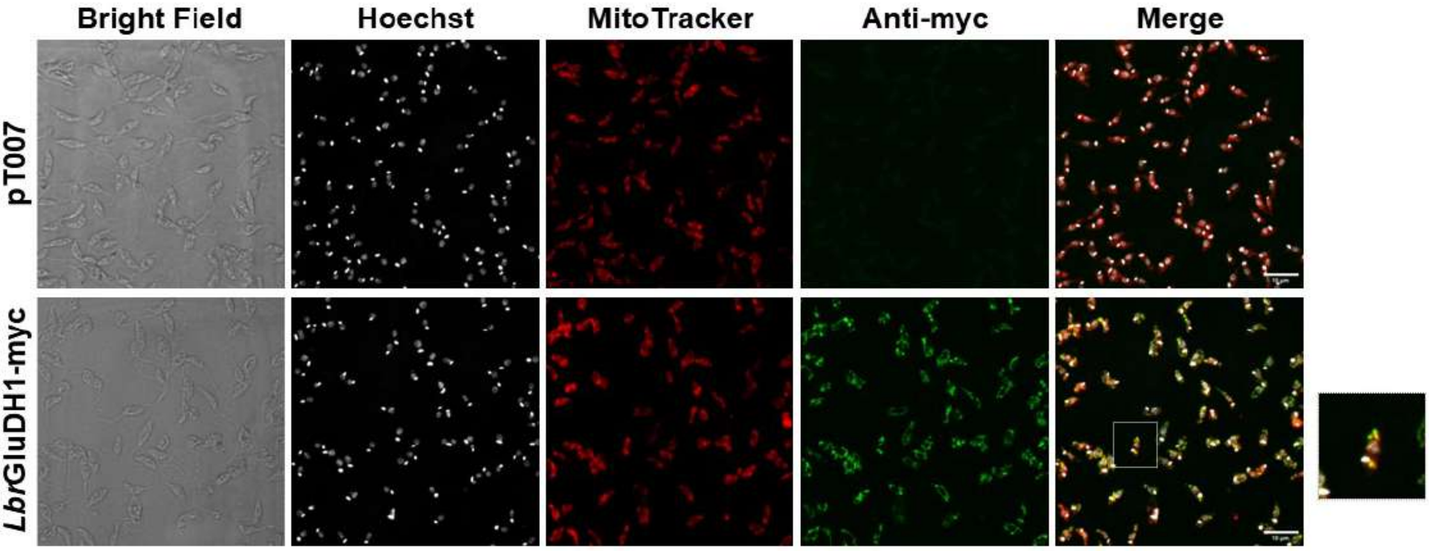


Supplementary Fig. 6

Supplementary Figure 6. *Lbr*GluDH1 localises to the mitochondria. The subcellular localisation of *Lbr*GluDH1-myc was determined by immunofluorescence against the myc tag. The the myc signal was completely superposed to the MitoTracker Deep Red mitochondrial dye.

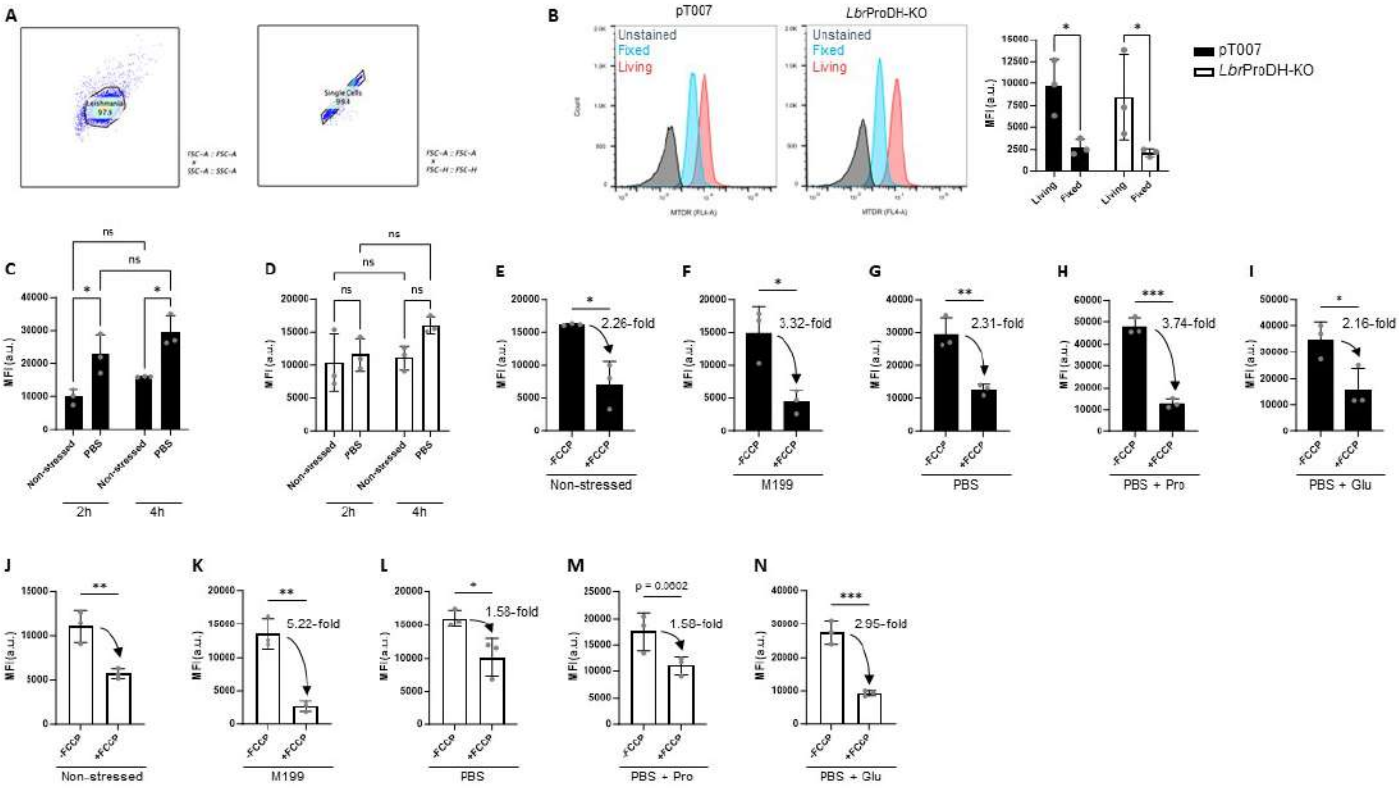


Supplementary Fig. 7

Supplementary Figure 7. Accumulation of MTDR depends on mitochondrial membrane potential in *L. braziliensis*. (A) Representation of the gating strategy used to isolate *Leishmania* single cells during analysis. (B) Living pT007 (black bars) and *Lbr*ProDH-KO (white bars) cells accumulated more MTDR than cells that were fixed in paraformaldehyde prior to incubation with MTDR: histograms displaying the fluorescence observed for unstained (grey), fixed (blue) and living (red) pT007 and *Lbr*ProDH-KO cells are shown on the left and on the centre, respectively; signal quantification in shown on the right. (C) pT007 cells subjected to acute nutritional stress accumulated more MTDR than cells kept in FBS-free M199, and this was unchanged from two to fours hours of stress. (D) *Lbr*ProDH-KO cells subjected to nutritional stress did not accumulate more MTDR than cells kept in FBS-free M199. (E – N) Addition of 10μM FCCP 30 min prior to the addition of MTDR during recovery of pT007 (black bars, E – I) and *Lbr*ProDH-KO (white bars, J – N) cells from nutritional stress significantly reduced MTDR accumulation in all conditions, except for *Lbr*ProDH-KO cells recovered in PBS + Pro (M, $p = 0.06$ by Student's t-test). (E and J) Non-stressed cells. (F – I and K – N) Cells stressed in PBS for two hours and transferred to M199 (F and K), PBS (G and L), PBS containing 3mM Pro (H and M) or 3mM Glu (I and N). *$P<0.05$; **$P<0.01$; ***$P<0.001$; ns – non-significant.

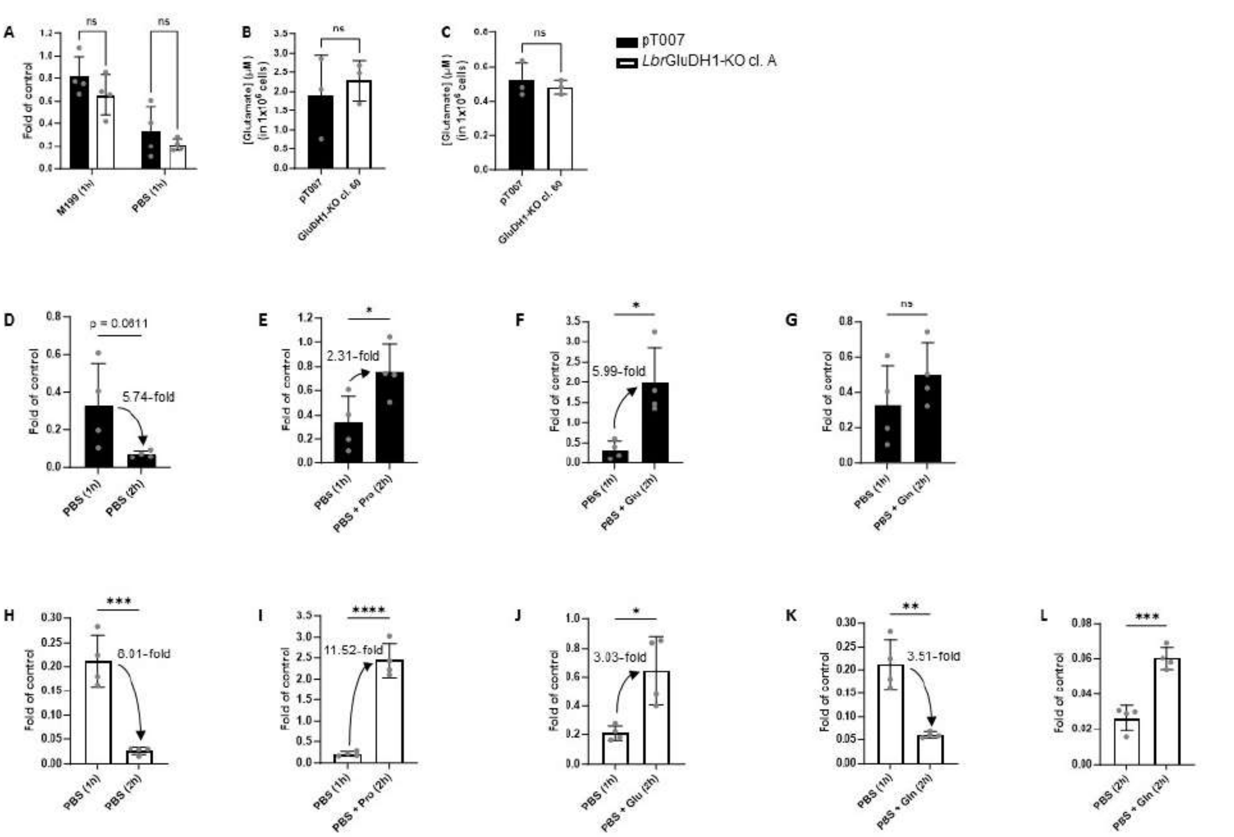


Supplementary Fig. 8

Supplementary Figure 8. Dynamics of the recovery of the intracellular levels of free Glu after nutritional stress and recovery. (A) pT007 (black bars) and *Lbr*GluDH1-KO (White bars) maintained similar proportional levels of intracellular Glu when kept for one hour in FBS-free M199, and the decayed in Glu levels was comparable between the two cell lines upon incubation in PBS for one hour. Incubation in PBS for one hour reduced the levels of intracellular Glu to similar levels in both cell lines (B). Both cell lines displayed similarly low levels of intracellular free Glu after incubation in PBS for two hours (C). pT007 (D – G) and *Lbr*GluDH1-KO (H – K) recovered their intracellular levels of free Glu in different fashions after one hour of starvation in PBS: (D and H) keeping the cells in PBS further reduced the levels of intracellular Glu, whereas the transference from PBS to PBS containing 3mM of Pro restored the levels of Glu in pT007 to close to what was observed in non-stressed cells at time 0 (E), but led to an accumulation of this amino acid in *Lbr*GluDH1-KO cells (I). Transference to PBS containing 3mM of Glu led to a two-fold accumulation of this amino acid in pT007 as compared to cells collected before nutritional stress (F). *Lbr*GluDH1-KO cells partially restored their levels of intracellular Glu when transferred to buffer containing 3mM Glu, but did not accumulate this amino acid within the cell (J). pT007 cells transferred to PBS + 3mM Gln neither restore their levels of intracellular Glu nor displayed decreased amounts of the amino acid (G); *Lbr*GluDH1-KO cells incubated with 3mM Gln still displayed a decrease in the levels of intracellular Glu as compared to cells stressed in PBS for one hour (K). This decrease in intracellular free Glu in *Lbr*GluDH1-KO in presence of Gln was, however, less pronounced than that observed in cells kept in PBS only (L). *P<0.05; **P<0.01; ***P<0.001; ****P<0.0001; ns – non-significant.

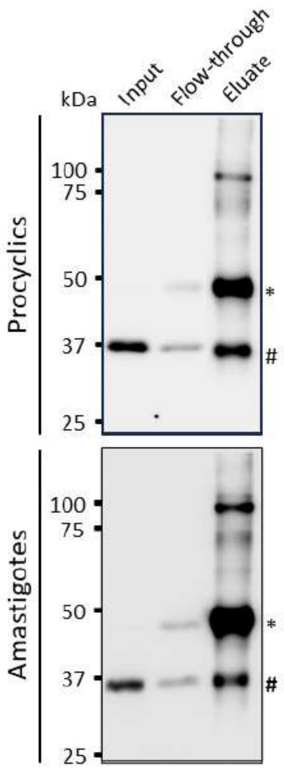


Supplementary Fig. 9

Supplementary Figure 9. Confirmation of myc-*Lbr*P5CR immunoprecipitation. The immunoprecipitation of myc-*Lbr*P5CR from procyclic (top) and amastigote (bottom) forms was confirmed by western blotting against the myc tag. *Anti-myc heavy chain; #myc-*Lbr*P5CR.
